# Hybrid E/M phenotype(s) and stemness: a mechanistic connection embedded in network topology

**DOI:** 10.1101/2020.10.18.341271

**Authors:** Satwik Pasani, Sarthak Sahoo, Mohit Kumar Jolly

**Author notes:** Authors to whom correspondence should be addressed (M.K.J.).

## Abstract

Metastasis remains an unsolved clinical challenge. Two crucial features of metastasizing cancer cells are a) their ability to dynamically move along the epithelial-hybrid-mesenchymal spectrum and b) their tumor-initiation potential or stemness. With increasing functional characterization of hybrid epithelial/mesenchymal (E/M) phenotypes along the spectrum, recent *in vitro* and *in vivo* studies have suggested an increasing association of hybrid E/M phenotypes with stemness. However, the mechanistic underpinnings enabling this association remain unclear. Here, we develop a mechanism-based mathematical modeling framework that interrogates the emergent nonlinear dynamics of the coupled network modules regulating E/M plasticity (miR-200/ZEB) and stemness (LIN28/let-7). Simulating the dynamics of this coupled network across a large ensemble of parameter sets, we observe that hybrid E/M phenotype(s) are more likely to acquire stemness relative to ‘pure’ epithelial or mesenchymal states. We also integrate multiple ‘phenotypic stability factors’ (PSFs) that have been shown to stabilize hybrid E/M phenotypes both *in silico* and *in vitro* – such as OVOL1/2, GRHL2, and NRF2 – with this network, and demonstrate that the enrichment of hybrid E/M phenotype(s) with stemness is largely conserved in the presence of these PSFs. Thus, our results offer mechanistic insights into recent experimental observations of hybrid E/M phenotype(s) being essential for tumor-initiation and highlight how this feature is embedded in the underlying topology of interconnected EMT and stemness networks.

## Introduction

Metastasis is the deadliest process in cancer, and it claims over 90% of all cancer-related deaths. It is a dynamic multi-stage cascade of events involving the initial detachment of cells from the primary tumor mass, dissemination of cancer cells, their exit (extravasation) at many distant organs, and finally, their ability to colonize distant organs through tumor outgrowth [1]. It is a challenging process for cells with extremely high attrition rates (> 99%) such that most cells die in circulation or are unable to adapt to foreign biochemical and/or biophysical ecology of distant organs [2,3]. Thus, phenotypic plasticity – the ability of disseminating cancer cells to adapt their phenotypes reversibly in response to their dynamic microenvironments – has been regarded as a hallmark of metastasis-initiating cells [4]. Phenotypic plasticity exists at multiple interconnected axes – a) epithelial-mesenchymal plasticity (EMP), b) metabolic plasticity, and c) plasticity between a cancer stem cell (CSC) and non-CSC state (i.e., stemness), among others [5]. Understanding such functional inter-dependencies from a dynamical systems level can lead to deciphering the organizing principles of underlying regulatory network modules and, eventually, therapeutic advances that can target cancer cell adaptability/plasticity [6,7].

Out of the various possible pairwise couplings among these different axes of plasticity, the one between EMP and stemness has been investigated most thoroughly through *in vitro, in vivo*, and *in silico* approaches. Initial reports, which treated EMP as a binary (all-or-none) process where cells switch between epithelial and mesenchymal phenotypes, suggested that a ‘full’ EMT (Epithelial-Mesenchymal Transition) was associated with increased stemness (tumor-initiation potential) [8,9]. This view was challenged by later studies demonstrating that a ‘full’ EMT was dispensable for acquiring stemness, or also perhaps an obstacle for the same [10,11]. A more nuanced recent view highlights that EMP involves multiple stable hybrid E/M states in addition to ‘fully’ E or ‘fully’ M ones and that these hybrid states may be maximally stem-like [12–17]. In squamous cell carcinoma cells that were categorized into six phenotypes along the EMP spectrum, stemness was shown to be acquired in earlier stages of EMP, and it did not increase as cells progressed towards a ‘fully’ mesenchymal phenotype. Intriguingly, the metastatic potential was found to be the maximum for hybrid E/M states [16]. Furthermore, in breast cancer cells, a transition from a hybrid E/M phenotype to a ‘fully’ mesenchymal one, as driven by constitutive ectopic expression of EMT-inducing transcription factor (EMT-TF) ZEB1, led to the loss of tumorigenicity *in vitro* and *in vivo* [13]. Three-dimensional assays to investigate collective cell invasion in breast cancer organoids showed that most leader cells co-expressed epithelial, mesenchymal, and CSC markers [18], further strengthening the association between hybrid E/M phenotype(s) and stemness. One possible reason for higher stemness of hybrid E/M cells may be their increased propensity to give rise to epithelial and mesenchymal cells [19,20] in addition to self-renewal, the two critical traits of stem cells in development and homeostasis. While this association of hybrid E/M phenotype with enhanced stemness has been consistently reported across cancer types [21], a mechanistic underpinning facilitating the association’s emergence remains unclear.

We have previously developed mechanism-based mathematical models that investigate the coupled nonlinear dynamics of EMP and stemness modules – miR-200/ZEB and LIN28/let-7, respectively. These models have predicted that while the most likely positioning of ‘stemness window’ is around the mid-point of ‘EMP axis’ (i.e., hybrid E/M phenotypes) [15], the ‘window’ is dynamic in nature and could move to either more epithelial or more mesenchymal ends too [22,23]. A caveat of this analysis is that it was performed on a few specific parameter sets; thus, its applicability to explain the diverse and mounting experimental evidence across cancer types, as mentioned above, remains limited.

Here, we have simulated the coupled dynamics of miR-200/ZEB, and LIN28/let-7 circuit over a wide range of biologically relevant parameter sets to interrogate whether the association of hybrid E/M phenotype with stemness is a trait embedded in the topology of this coupling itself. Our results suggest that the overlap of hybrid E/M and stemness features is an outcome of coupled network topology between the miR-200/ZEB and LIN28/let-7 feedback loops. Moreover, upon extending the network to include various phenotypic stability factors (PSFs) such as GRHL2, OVOL1/2, and NRF2, we notice that this association is maintained. Thus, our results unravel the underlying design principles of EMP-stemness association and predict that this interconnection is likely to be seen across multiple carcinomas.

## Results

### Gene regulatory network underlying E/M plasticity and stemness enables the existence of four E/M phenotypes

To unravel the mechanistic underpinnings of the association between stemness and epithelial-mesenchymal plasticity (EMP), we considered the regulatory interactions among key factors implicated in governing the E/M plasticity (ZEB/miR-200) and the stemness characteristics (LIN28/let-7) **(Fig 1A)**. E/M plasticity is at least in part controlled by a mutually inhibitory feedback loop between the transcription factor family ZEB and the microRNA family miR-200 [24,25]. Moreover, ZEB can self-activate through ESRP1 and/or CD44/HA signaling [26–28]. This network can give rise to three phenotypes: epithelial (high miR-200, low ZEB), mesenchymal (low miR-200, high ZEB), and hybrid E/M (medium miR-200, medium ZEB) [29]. This network can be influenced by other EMT-TFs such as SNAIL which self-inhibits transcriptionally and activates the expression of ZEB, and suppresses the transcription of the microRNA miR-200 [29]. Similarly, the ‘stemness’ of a cell is primarily controlled by a mutually inhibitory feedback loop between RNA-binding factor family LIN28 and microRNA family let-7 [30]. Both LIN28 and let-7 are known to self-activate through direct and/or indirect mechanisms [31–33], and the transcription factor NF-kB activates the expression of both of them [15]. LIN28 regulates the levels of OCT4 [34], whose intermediate levels are considered to be optimal for attaining stemness [35–37]. Interestingly, these two modules – ZEB/miR-200 and LIN28/let-7 – can influence each other, such that let-7 can target ZEB translation [33], and microRNA family miR-200 can inhibit LIN28 translation [38,39]. Put together, this integrated network of EMP and stemness regulators can explain experimental observations on LIN28 driving EMP [40] and ZEB/miR-200 influencing stemness [41].

**Figure 1:**
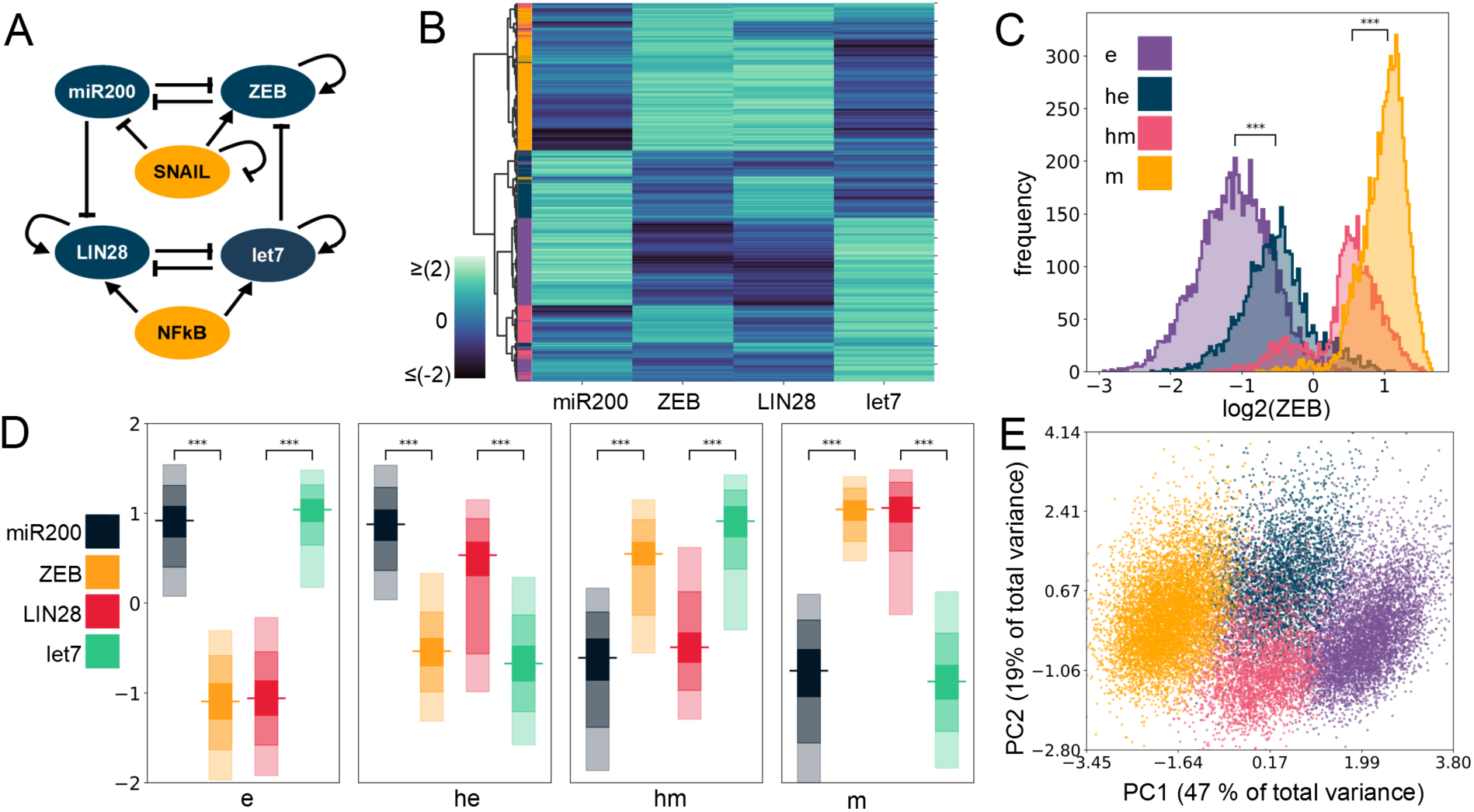
Four E/M phenotypes enabled by the coupled EMP-stemness circuit. **A)** The core EMP-stemness circuit (base circuit). Arrows represent activating links and hammerheads represent inhibitory links. **B)** Heatmap showing the steady state expression of all solutions of a single replicate including the core EMP (ZEB, miR-200) and stemness (LIN28, let-7) nodes. Expression histograms (all solutions from a single RACIPE replicate. For n=5, see **Supplementary Table S2** in **section 2.1**) showing ZEB levels in the 4 clusters – e, he, hm and m. The colors of the labels shown in heatmap are for e, he, hm, m clusters displayed here. Expression levels of EMP and the stemness genes in the 4 clusters. In each node, the horizontal line represents the median value, the darkest region represents the middle 30% (35^th^ to 65^th^ percentile) data, the region next in lightness represents the next 40% (15^th^ to 35^th^ and 65^th^ to 85^th^ percentiles), while the lightest region represents the next 20% of the data (5^th^ to 15^th^ and 85^th^ to 95^th^ percentiles) **E)** PCA plot colored according to the K-Means cluster assignment. (PC1 = 0.48 x miR200 – 0.54 * ZEB – 0.21 x SNAIL – 0.49 x LIN28 + 0.44 x let7 + 0.057 x NF-kB; PC2 = 0.27 x miR200 – 0.08 x ZEB – 0.66 x SNAIL + 0.14 x LIN28 – 0.49 x let7 – 0.47 x NF-kB). For C, D: Significance bars added to show statistically significant differences (Mann-Whitney U test). Legend for the p-values +: p > 0.01, *: 0.01 > p > 0.001, **: 0.001 > p > 0.0001, ***: p < 0.0001; for effect sizes refer to **Table S7** in **Supplementary Section 2.5**

All these regulatory links in the abovementioned network may be active with different strengths in individual cells. Further, in cancer cells, the mutational background can alter the stability and/or production rate of various species, for instance, mutant p53 gain of function, triggering ZEB1 [42]. Thus, a rigorous analysis of the emergent dynamics of the coupled EMP-stemness network would require a framework that can decode the features robust to parameter variation of specific network topology. To achieve this goal, we simulated the coupled EMP-stemness network through RACIPE (Random CIrcuit Perturbation) [43] (see **Methods**). RACIPE takes network topology as its input, converts it into a set of coupled ordinary differential equations (ODEs) where the influence of each activation or inhibition regulatory link is represented by a shifted Hill function. Parameters corresponding to this set of equations are sampled randomly within a biologically relevant range. Therefore, it generates an ensemble of ODE models, each with a different parameter set, thus representing the intrinsic variability in kinetic parameter values in a given cell population. The set of steady state values obtained from this ensemble can be then used to identify robust dynamical signatures emerging from this topology through appropriate statistical analysis (see **Methods**).

First, we plotted a heatmap of all the steady-state values obtained via RACIPE for this coupled network and performed agglomerative hierarchical clustering **(Fig 1B)** on it. We observed the possible existence of four major clusters, indicated in the dendrogram plotted using Ward’s minimum variance criterion (see **Methods**) **(Fig S1A)**. We next performed K-means clustering for varied values of K and observed a peak in the average silhouette width at K=4, endorsing the existence of four clusters in RACIPE solutions **(Fig S1B)**. This trend was supported by all other cluster quality metrics we used (**Fig S1C-E)**. We plotted histograms of ZEB levels for the four clusters identified in the heatmap and assigned them phenotypic labels accordingly **(Fig 1C, 1D)**. The cluster with the minimum median ZEB levels is referred to as epithelial (e) phenotype, while the cluster with the maximum ZEB median levels is referred to as a mesenchymal (m) phenotype. The other two clusters had intermediate ZEB levels being categorized as E/M hybrids and are assigned the labels of hybrid epithelial (he) and hybrid mesenchymal (hm). Furthermore, we performed principal component analysis (PCA) on all steady-state solutions **(Fig 1E**; each color represents a distinct cluster identified by K-means). Principal Component 1 and 2 explain 47% and 19% of the total variance, respectively, and show roughly 4 clusters in the PCA plot where the two hybrid clusters (blue and pink) are sandwiched between the epithelial (purple) and the mesenchymal (yellow) clusters (**Fig 1E**). PC1 can be thought of as an approximate EMT axis based on the coefficients for miR-200 and ZEB. Interestingly, while miR-200 levels are very different between e and m states, its levels are similar for e and he clusters, and for hm and m clusters **(Fig S2A, 1D, Table S7)**, thus indicating that miR-200 may be a more coarse-grained readout of EMT state of a cell. Corroborative trends, i.e. highest levels of let-7 and lowest levels of LIN28 in the e cluster, and lowest levels of let-7 and highest levels of LIN28 in the m cluster were observed (**Fig 1D, S2B**). The levels of NF-kB and SNAIL levels were found to be almost similar across the clusters **(Fig S2C-D)**, which is not surprising because neither NF-kB nor SNAIL are influenced by any of the other four nodes (miR-200, ZEB, let-7, LIN28) in the coupled EMP-stemness network.

### Stemness characteristics are enriched for in the hybrid E/M phenotypes

The gene regulatory network involving the interplay between E/M plasticity and stemness can give rise to different phenotypes (steady-states). Next, we investigated whether these states can co-exist for certain parameter sets, i.e. is the network capable of exhibiting multistability? Out of the ensemble of parameter sets sampled by RACIPE, we observed that a subset of parameter sets can give rise to a combination of two or more steady-states per parameter set (bistability: 37.3 +/- 0.95%, tristability: 24.2 +/- 0.09%, etc.) while the others converged to a single steady-state for the given system of ordinary differential equations (monostability: 29.6 +/- 0.70%). A closer inspection of the monostable solutions revealed epithelial (e) and mesenchymal (m) states as the most predominant solutions **(Fig 2A)**, indicating that epithelial and mesenchymal phenotypes are more likely to exist relative to the hybrid ones, thereby reminiscent of observations based on modeling of larger EMP regulatory networks too [44,45]. This trend is also observed when considering all the solutions together irrespective of the number of states (**Table S5**). Consistently, among all possible bistable solutions, the most predominant one is the phase {e, m}, i.e. the co-existence of epithelial and mesenchymal states **(Fig S3A)**. The least frequent bistable phase is {he, hm}, i.e. the one with co-existence of both hybrid phenotypes; while the frequency of bistable phases containing one hybrid phenotype and one of the epithelial or mesenchymal ones – {e, hm}, {he, m}, {e, he}, {m, hm} – is intermediate **(Fig S3A)**. Similarly, among all possible tristable solutions, the phases containing both epithelial and mesenchymal states - {e, he, m} and {e, hm, m} – were the most predominant **(Fig S3B)**. Put together, these results indicate that the coupled EMP-stemness regulatory network is multistable and therefore allows for cells to spontaneously switch among the epithelial, mesenchymal, and the hybrid phenotypes.

**Figure 2:**
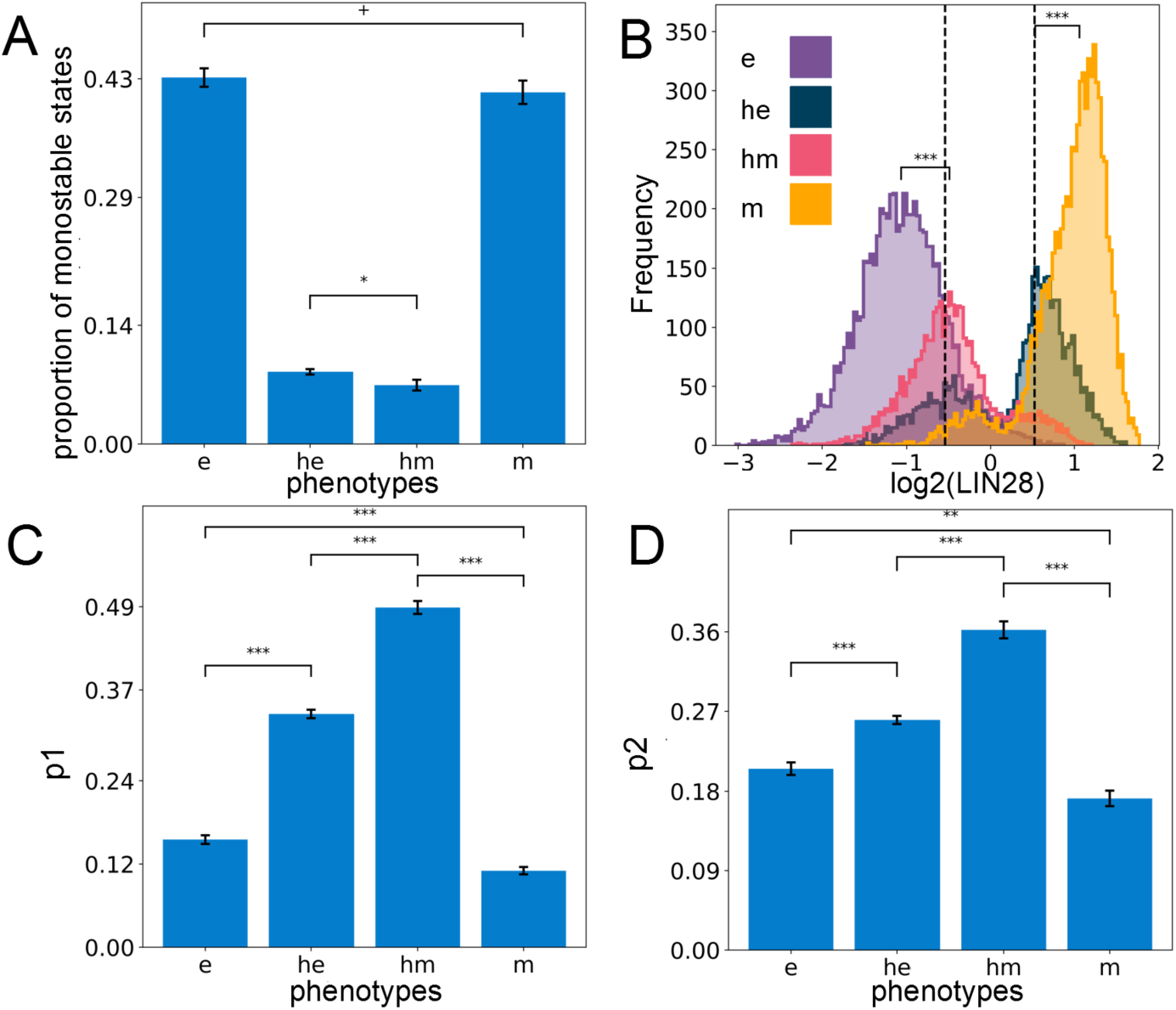
Hybrid E/M phenotypes are enriched for stemness. **A)** Proportion of all monostable parameter sets belonging to different phenotypes – e, he, hm, m. **B)** Expression histogram for LIN28 (all solutions of a single RACIPE replicate) for the four phenotypes with the stemness window (black vertical lines) as defined by the middle 30% of the ‘LIN28 biological range’ (median ± one inter-quartile range averaged over n=5 RACIPE replicates). Significance bars added to show statistically significant differences (Mann-Whitney U test) **C)** Value of p1 - the probability of a solution lying in the stemness window, conditional on the solution belonging to a particular phenotype for all the phenotypes (n=5 RACIPE replicates). **D)** Same as C) but for p2 - the probability of a solution belonging to a particular phenotype conditional on it lying in the stemness window. For A, C, D: mean values show average across five RACIPE replicates and the error bars represent the across-replicate standard deviations. Two-tailed Students’ t-test with unequal variance (Welch’s t-test) was performed. Legend for the p-values +: p > 0.01, *: 0.01 > p > 0.001, **: 0.001 > p > 0.0001, ***: p < 0.0001; for effect sizes refer to **Table S7** in **Supplementary Section 2.5**.

Next, we investigated the stemness traits of different cell states along the E/M spectrum. Intermediate levels of OCT4 have been reported to confer maximal stemness to the cells [35–37]. Given that OCT4 is a direct target of LIN28, we defined the “stemness window” to be a region around median LIN28 expression levels in all steady-state solutions combined and the median averaged over all the replicates of the coupled EMP-stemness circuit. Specifically, we considered the median ± one inter-quartile range of the LIN28 distribution to be the “biological range” of LIN28, and the middle 30% of this range centered around its median is considered to be the ‘stemness window,’ which might allow for a population enriched in the stem-like phenotype **(Fig 2B)**. To test the robustness of this specifically chosen stemness window, we need a semi-objective heuristic for increasingly wider “stemness windows” centered at the mid-point of the range chosen. When the size of the ‘stemness window’ is considered to be more than one-third of the range of LIN28 levels, we see a qualitative change in the distribution of phenotypes lying in this window. This analysis underscores that hybrid E/M phenotypes are predominantly associated with stemness unless the ‘stemness window’ starts getting too large to be of a biologically implausible size **(Fig S3C)**.

Next, we quantified the probability that a particular solution lies within the “stemness window” given it belongs to a particular phenotype along the E-M spectrum (p1). Interestingly, we found that hybrid (he and hm) phenotypes are more likely to be stem-like than either extreme (e and m) E/M phenotypes **(Fig 2C)**. Alternatively, we computed the probability that a solution belongs to a particular phenotype along the E-M spectrum, given that it already lies in the “stemness window” (p2). Congruently, we observed that hybrid phenotypes (he and hm) are enriched for in the stemness region than either epithelial (e) or mesenchymal (m) solutions, although the contribution of e and m cell states to “stemness window” cannot be ignored **(Fig 2D)**. Put together; these observations suggest that although the hybrid phenotypes he and hm are more likely to be associated with a stem-like characteristic, stemness is not exclusive to them. Thus, a subset of ‘pure’ epithelial and/or mesenchymal cell states can also be potentially stem-like, depending on parametric combinations.

Having established a strong association between the hybrid E/M phenotypes and stemness, we sought to understand the mechanistic underpinnings that can facilitate this association. Through RACIPE, we simulated a hypothetical ‘uncoupled’ circuit where the inhibitory links from miR-200 to LIN28 and that from let-7 to ZEB are absent. The ensemble of solutions obtained for this uncoupled circuit also has four clusters (see **Supplementary Section 2.2**). For the uncoupled circuits, when we plot the clusters in a ZEB-LIN28 plane (indicative of an EMP-stemness plane with ZEB being a proxy for E/M axis and LIN28 as a proxy for stemness), we observed that the mid-points of these four clusters were arranged in a square – the four vertices being [high ZEB, high LIN28], [low ZEB, high LIN28], [high ZEB, low LIN28] and [low ZEB, low LIN28] – demonstrating the independence of the ZEB and LIN28 levels **(Fig S4A)**. The [high ZEB, high LIN28] and [low ZEB, low LIN28] clusters correspond to m and e states respectively and the location of the former on the two-dimensional plane for the coupled circuit overlapped with that of the uncoupled circuit. However, in case of the coupled circuit, the square geometry is disturbed; instead, more of a rhombus-like shape emerged due to the shift of the mid-points of the [low ZEB, high LIN28] and [high ZEB, low LIN28] clusters which now tend to have intermediate values of both. This analysis suggests that coupling EMP and stemness modules enables a higher likelihood of both hybrid (he, hm) clusters to be more stem-like **(Fig S4A)**. This observation can be mechanistically interpreted by understanding the relative ‘activity’ of the two coupling links (miR-200 inhibiting LIN28 and let-7 inhibiting ZEB). In [high ZEB, high LIN28] state, the levels of let-7 and miR-200 are relatively low, thus, the links are effectively ‘inactive’. In case of [low ZEB, high LIN28] and [high ZEB, low LIN28], one link is much more active than the other, hence we see relocations mostly along one of the two axes of the LIN28-ZEB plane. For the [low LIN28, low ZEB] state, both links are approximately equally ‘active’, and therefore the cluster migrates along to even lower values of both LIN28 and ZEB.

To validate this hypothesis, we quantified the strength of coupling links – the net effect was strongest for e (i.e. [low LIN28, low ZEB]) and weakest for m (i.e. [high ZEB, high LIN28]) (**Fig S4B**). As a control case for the link strength metric, we quantified the degree of asymmetry in the strength of the two links within EMP and stemness modules individually (See **methods)**. We observed that as expected, the effective “suppression” (the expression free topological asymmetry) of ZEB on miR-200 was the highest in the case of hm and m states relative to e and he states **(Fig S4D)**. On the other hand, we observed that the suppression of LIN28 on let-7 was higher in the case of he and m and lower in the e and hm phenotypes **(Fig S4C)**.

### The effect of phenotypic stability factors (PSFs) on the phenotypic landscape of EMP

Previous work, including ours, has shown that various factors, such as GRHL2, OVOL1/2, and NRF2, can stabilize the hybrid E/M phenotype of cells [46–49]. Their knockdown in H1975 – a stable hybrid E/M lung cancer *in vitro* model – pushed cells towards a complete EMT, thus, they were classified as ‘phenotypic stability factors’ (PSFs) for hybrid E/M phenotypes. For a specific set of parameters, incorporating GRHL2 also promoted the association between hybrid E/M states and stemness [46]. Thus, we investigated whether the hybrid E/M-stemness interconnection was maintained for multiple PSFs across an ensemble of parameter sets.

We simulated via RACIPE the coupled EMP-stemness networks after incorporating these three PSFs, one at a time **(Fig 3A)**. As compared to the circuit without these PSFs (i.e., the ‘base’ coupled EMP-stemness network shown in Fig 1A), the circuits including PSFs had a higher frequency of parameter sets leading to tristability and quadrastability, at the expense of lower frequency of parameter sets leading to monostability primarily and even bistability in the case of the GRHL2 circuit (**Fig 3B**). In general, all these circuits showed a trend towards increasing multistability, with the strongest effect seen for the GRHL2 circuit **(Fig 3B)**. However, incorporating the PSFs altered neither the optimal number of clusters as identified by average silhouette widths **(Fig 3C)** nor the position of clusters on the ZEB-LIN28 plane that showed an almost complete overlap with those seen for the ‘base’ circuit (**Fig 3D**). Therefore, the introduction of the PSFs does not seem to disrupt the core structure of the steady-state solutions as obtained earlier.

**Figure 3:**
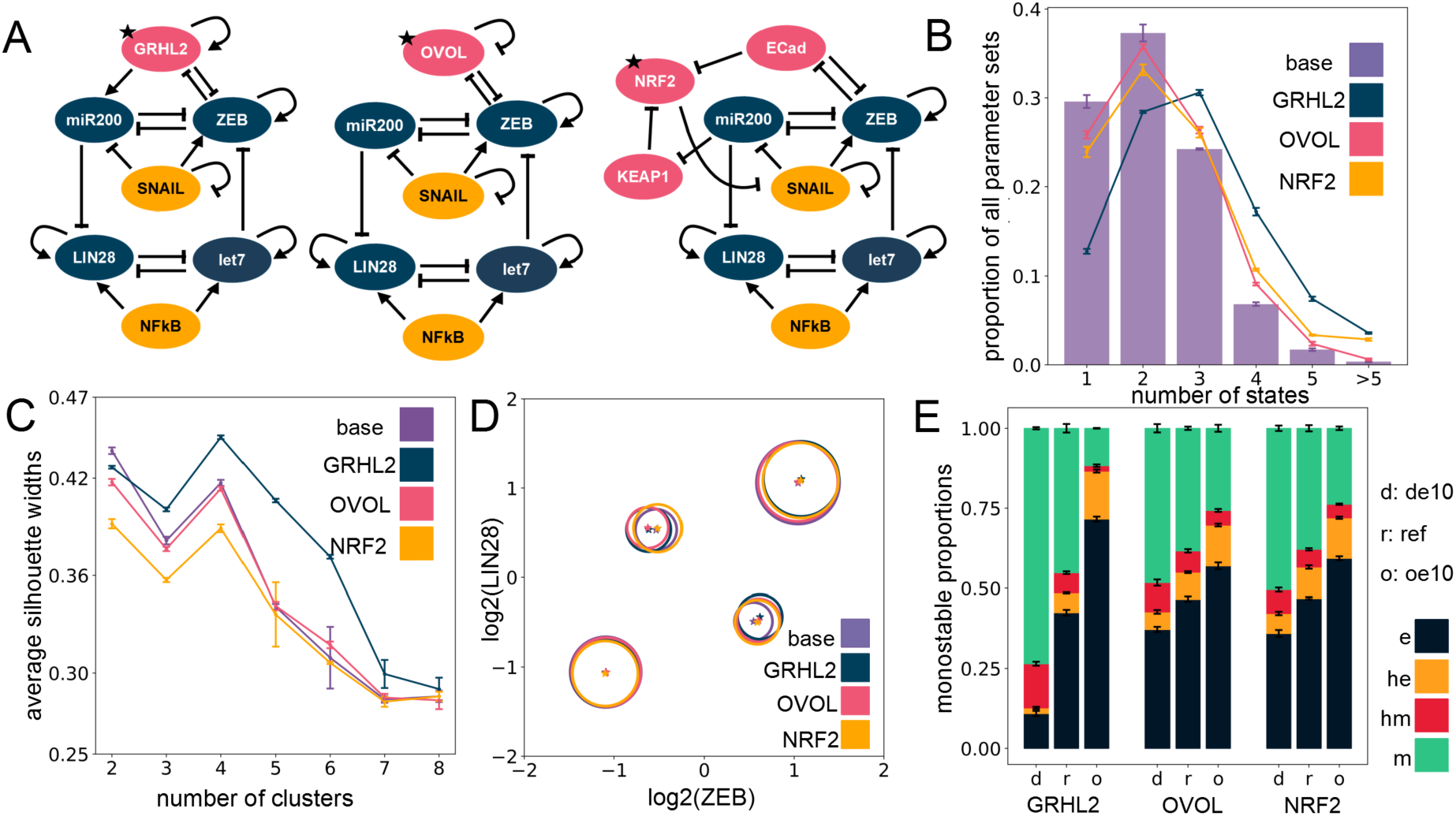
Including PSFs preserves the cluster structure seen for EMP-stemness circuit. **A)** Different PSFs (GRHL2, OVOL2, NRF2) added to the base (control) EMP-stemness circuit. All new nodes are shown in pink; the PSF node added is marked with a star. **B)** Multi-stability profiles for all the PSF circuits compared to the base circuit. **C)** The average silhouette widths at different numbers of clusters for all PSF circuits with the cluster membership assignment via K-Means algorithm. Note the distinct peak at cluster number = 4 across all circuits. **D)** Cluster centers in the ZEB-LIN28 plane (EMP-stemness plane) for a single RACIPE replicate of each of the circuits show that the cluster centers are well-preserved across the circuits. Gene-wise normalization is performed independently for all circuits (see **Supplementary section 1.2.3**). **E)** Phenotypic distributions for all monostable solutions upon overexpressing and down-expressing the different PSF nodes. (de10: 10-fold downexpression, ref: reference unperturbed circuit, oe10: 10-fold overexpression) (B, C, E: The values are averaged across n=5 RACIPE replicates and the error bars represent the across-replicate standard deviations)

In addition to being thought of as PSFs, GRHL2 and OVOL1/2 have been considered as MET-inducing transcription factors (MET-TFs) as their overexpression in carcinomas is capable of upregulating the expression levels of E-cadherin and/or revert EMT [50–54] as well as other EMT-associated traits such as anoikis-resistance, metabolic reprogramming [55] and immune evasion [56]. This observation is consistent with their knockdown known to drive a ‘full’ EMT in both cancer cell lines and in developmental contexts [46,49]. Thus, with the goal to identify the effect of overexpression and downregulation of these PSFs on relative proportions of the different EMP, we quantified the changes in the frequency of different monostable solutions – e, m, he and hm – upon a 10-fold overexpression (oe10) and a 10-fold down-expression (de10) of these PSFs. Computationally, this provides a dynamically consistent way to characterize the effect attributable to any node on the network dynamics. and RACIPE implements this perturbation by altering rate of production of the specific node (see **methods**). We observed that increasing the expression of any one of the three PSFs enriched the epithelial (e) phenotype at the expense of mesenchymal (m) phenotype. Similar, albeit weaker, trends were seen for an enrichment of the he phenotype at the expense of the hm phenotype **(Fig 3E)**. This effect was more pronounced in the GRHL2 case in comparison to either OVOL or NRF2 **(Fig 3E)**. This trend is also present overall not only for only monostable solutions, but also for all parameter sets considered together (see **Supplementary section 2.4**).

Moreover, a closer inspection of steady-state solutions from bistable parameter sets showed a consistent trend across PSFs in case of their overexpression: most remarkable decrease in the proportion of {hm, m} phase with the most remarkable increase in the proportion of the {he, e} phase **(Fig S5A-C)**. Similarly, the tristable states also showed an agreement with the trend of enriching “relatively epithelial” phases, with maximum enrichment of the {e, he, m} phase and a maximum depletion of {e, hm, m} phase on overexpression of the PSFs **(Fig S5D-S5F)** with unremarkable changes in the other phases (see **Supplementary Section 2.3** for details on the multistable distributions). Put together; these results suggest that incorporating any of these PSFs (GRHL2, OVOL, NRF2) tends to maintain the 4 clusters seen in EMP-stemness coupled network earlier, and their overexpression can serve as a way to inhibit at least the progression of a complete EMT.

**Networks incorporating Phenotype Stability Factors (PSFs) preserve the association of the hybrid E/M states with stemness**

Next, we evaluated how the association of EMP phenotypes with stemness is altered upon including PSFs. As earlier, we explored two kinds of association of the hybrid E/M phenotypes and a stem-like state: the probability of being a stem-like state given that a solution belongs to a particular E/M phenotype (p1) and the probability of belonging to a particular E/M phenotype given that the solution is a stem-like state (p2). On comparing p1 and p2 for all circuits with the two hybrid clusters (he, hm) taken together and the two non-hybrid clusters (e, m) taken together, we observe that the hybrids are much more likely to be stem-like **(Fig 4A)** as well as constitute a larger proportion of the stemness enriched pool of solutions **(Fig 4B)**, across all cases of PSFs. This observation suggests that the association of hybrid E/M phenotypes with stemness is a dynamical trait encoded in the core network topology, and this feature is maintained upon including PSFs.

**Figure 4:**
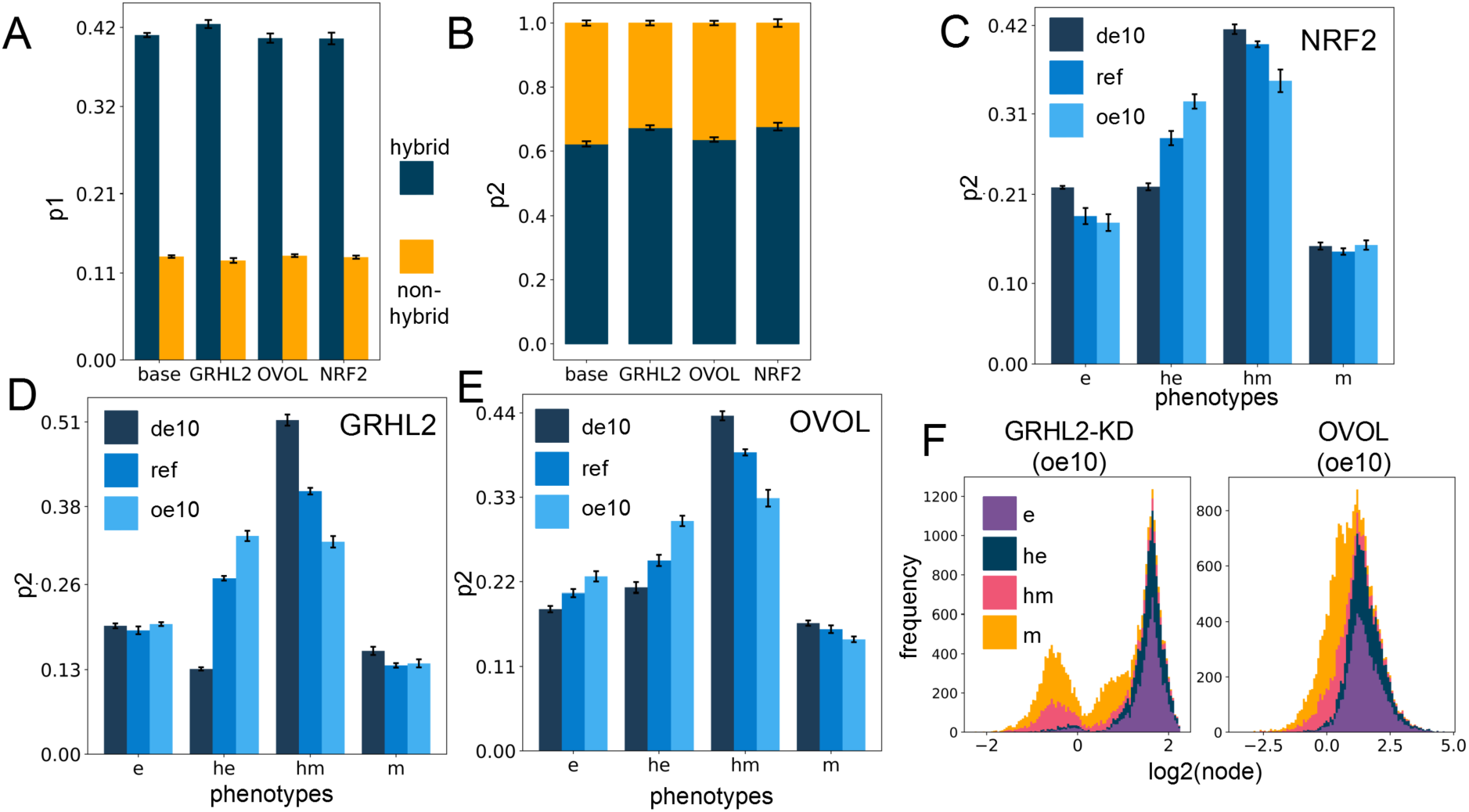
PSF motifs maintain the association between hybrid E/M states and stemness. **A)** Sum of p1 values for the two hybrid (he, hm) and two non-hybrid (e,m) phenotypes, for different circuits including PSFs. **B)** Same as A) but for p2. **C-E)** p2 values for four phenotypes for cases of overexpression and downregulation of each of the PSFs individually: NRF2 (C), GRHL2 (D) and OVOL (E). **F)** Stacked expression histograms (colored in accordance with the phenotypes) showing the change in the PSF node distribution in the GRHL2-KD circuit (GRHL2-KD) (left panel) vs the OVOL circuit (right panel) at 10-fold overexpression. GRHL2-KD involves knockdown of the activating link from GRHL2 to miR-200, causing the only difference between the two presented circuit to be the nature of the self-link of the PSF node. All other links remain the same. (de10: 10-fold downexpression, ref: reference unperturbed circuit, oe10: 10-fold overexpression) (A-B, C-E: The values are averaged across replicates and the error bars represent the across-replicate standard deviations)(refer **Table S8** in **Supplementary Section 2**.5 for the statistical testing of these quantities)

A closer look at the changes in p1 and p2 values upon overexpression or downregulation of PSFs revealed small variable minor changes for p1 **(Fig S6A-C)**, but the p2 values for hybrid epithelial (he) phenotype increased whereas that of hybrid mesenchymal (hm) state decreased with little changes in that of e or m phenotypes **(Fig 4C-E)**. However, at all levels of PSF expression in all circuits (from 10-fold down-expression to a 10-fold overexpression), the hybrids together (he, hm) have a more considerable value of p1 and p2 in comparison to the ‘fully’ epithelial and mesenchymal states (e, m) **(Fig 4C-E, S6A-C)**, suggesting that hybrid E/M phenotypes are more likely to be stem-like than either extreme of the EMT axis. Further, over-expression of PSFs may pull the composition of the solutions in the ‘stemness window’ towards the epithelial end of EMT axis.

We additionally looked at the distribution profile of the PSFs in all the clusters **(Fig S7A-I)** for all the circuits with down- and over-expression. Overall, the distribution profiles are distinct for all the circuits, perhaps indicating the role of specific topological features in dictating this distribution profile. GRHL2 remains appreciably bimodal throughout the range of expression, and the e and he clusters occupy the higher end of the GRHL2 histogram, while the m and hm clusters occupy the lower end **(Fig S7A-C)**. OVOL is bimodal at 10-fold down-expression, but the profile shifts towards unimodality on increasing expression, possibly due to its self-inhibition. Generally, a lower OVOL level is associated with the m and hm clusters and vice versa **(Fig S7D-F)**. On the contrary, NRF2 distribution profile does not show any readily appreciable preference for any of the clusters at any level of expression **(Fig S7G-I)**.

The preference of GRHL2 and OVOL for e and he cluster may be attributed to the presence of a mutually inhibitory negative feedback loop with ZEB, which causes the PSF (GRHL2, OVOL) levels to be relatively high when the ZEB levels are low (e and he) and *vice versa*. Tge GRHL2 and OVOL coupling with EMP circuit has two differences: a) GRHL2 activates miR-200 but OVOL does not, and b) GRHL2 self-activates but OVOL self-inhibits. To deconvolute the effect of these two differences, we investigated a circuit where the activation link from GRHL2 to miR-200 is knocked down (GRHL2-KD). This circuit differs from the OVOL circuit only in terms of the nature of the self-regulation. We then investigated this knockdown circuit in regard to all the investigations explored for each of the other circuits above and compared them to the trends observed for the OVOL circuit. Qualitatively, all the trends matched well, and the only remarkable difference was in the cluster-wise distribution profile of the PSF at different levels of expression **(Fig 4F, S8A-F)**, suggesting that the specific enrichments observed are primarily dictated by the interacting links between the PSF node and the other nodes, while the PSF distribution profile itself is strongly affected by the nature of the self-link.

## Discussion

Metastasis has been indicated as a hallmark of cancer [57]. However, decades of research have highlighted that hallmarks of metastasis are quite different from those of primary tumor growth [58]. Given the ever-changing biochemical and biomechanical environments for the cell in the metastatic cascade, it is not surprising that metastasis is a highly inefficient process and that only a minuscule percentage of disseminated cells that can pass through these bottlenecks by adapting to their dynamic environments can eventually colonize distant organs and form macrometastases. Thus, the concept of Lamarckian evolution seems to be more prevalent than that of Darwinian selection (of a subset of genetic clones) in the context of metastasis. Consequently, environment-driven phenotypic plasticity has been advocated as a hallmark of metastasis [58]. Recent observations in the evolution of cancer therapy resistance endorse the role of Lamarckian induction in enabling the dynamic adaptability of cancer cells [59–63].

Here, we elucidate the design principles of regulatory networks connecting two crucial aspects of phenotypic plasticity during metastasis – epithelial-mesenchymal plasticity (EMP) and stemness. Our results demonstrate that a higher likelihood of hybrid epithelial/mesenchymal (E/M) phenotype(s) acquiring increased stemness is an outcome of the network topology of coupled EMP-stemness network. These findings offer a mechanistic basis of various recent *in vitro, in vivo*, and clinical work suggesting an enhanced stemness and/or aggressiveness trait of hybrid E/M phenotype(s), as reported across different carcinomas. A recent quantitative analysis of about a hundred urothelial carcinomas concluded that “it is not the amount, but merely the presence of a minimum of tumor cells in hybrid E/M states that contribute to disease aggressiveness” [64]. This clinical observation is reminiscent of *in vivo* experiments showing that both ‘extremely epithelial’ (generated via CRISPR/Cas9 by eliminating Zeb1 expression) and ‘extremely mesenchymal’ cells (generated by constitutive overexpression of Zeb1), either alone or in combination, had minimal tumor-initiation potential as compared to hybrid E/M cells. Put together, these results highlight the importance of individual cells co-expressing epithelial and mesenchymal markers (i.e. one or more hybrid E/M phenotype(s)) in enabling stemness. At least two factors can underlie this enhanced stemness of hybrid E/M cells: a) their ability to self-renew, as some cells undergoing a full EMT can undergo cell cycle arrest [65,66], and b) their ability to give rise to heterogeneous subpopulations of epithelial and mesenchymal cells [19,20] that can potentially cooperate in driving metastatic traits [67]. Thus, hybrid E/M cells can be conceptually thought of as similar to adult stem cells in a tissue, a notion supported by a hybrid E/M phenotype seen in a subset of mammary stem cells [68,69].

It should be noted that we are proposing an enrichment of hybrid E/M phenotype(s) in stem-like populations, but not an exclusive or exhaustive association between hybrid E/M cells and stemness. Neither every cell undergoing EMP, even a partial EMT, is expected to be stem-like; nor every cancer stem cell (CSC) is expected to manifest the hybrid E/M phenotype(s). CSCs of varying EMP status have been seen in breast cancer [11], prostate cancer [70], cervical cancer [71], and squamous cell carcinoma [72,73], with different possible spatial localization within a tumor [74]. This possibility can be explained by our model predictions demonstrating the presence of more epithelial or more mesenchymal phenotypes within the dynamic “stemness window,” albeit at a relatively lower frequency as compared to that of hybrid E/M phenotype(s).

Our results show that stemness as a function of EMP is not necessarily always monotonically increasing; similar observations have been reported now for tumor aggressiveness as a function of chromosomal instability (CIN) [75,76]. Such reports caution us against falling prey to tacit assumptions about monotonic relationships between any two molecular factors and/or phenomena and reveal the fundamental insights that can be gained by mechanistic modeling of nonlinear dynamics embedded in a complex regulatory network.

To investigate the robustness of our results and quantify the influence of various PSFs on the association between hybrid E/M phenotypes and stemness, we incorporated the PSFs GRHL2, OVOL2, and NRF2 in our coupled network. We observed that these PSFs maintained the enrichment of hybrid E/M phenotypes in stem-like traits, endorsing the robustness of this association. Consistently, many of these PSFs have been reported to play a functional role in maintaining more epithelial phenotypes (e, he) as well as stemness in experiments in cancer cell lines as well as in cellular reprogramming contexts involving modulation of EMP [53,77,78]. GRHL2 and OVOL2 have also been proposed as MET-inducing transcription factors (MET-TFs), but our simulations suggest that their ability to induce a ‘complete’ MET need not be universal. This prediction is validated by recent experimental observations that the ability of GRHL2 to induce a MET depended on the epigenetic status of specific cell lines where it is overexpressed ectopically [54,79]. Thus, our results propose that similar to variations in the ability of EMT-TFs to drive an EMT [80,81], MET-TFs may induce MET to varying degrees.

The approach presented here can be expanded to answer a long-standing open question in metastasis – how do cells coordinate various axes of their plasticity during the metastatic cascade? Mechanism-based models of individual axes of plasticity are the building blocks required to answer this question from a dynamical systems perspective. There has been increasing interest in dynamical modeling of EMP [82–86]; similar efforts to model related aspects of EMP such as metabolic reprogramming [87], autophagy [88,89], therapy resistance [90,91], and immune evasion [92] are being attempted too. Coupling these modules to decode the interconnections among different axes of plasticity can help unravel the survival strategies of metastasis-initiating cells and eventually contribute to designing new clinical strategies. For instance, a recent prediction we made is that breaking the positive feedback loops in a network can restrict plasticity and potentially impact metastatic potential [6]. Indeed, proof-of-principle validation for this prediction was seen when cancer cells with a compromised ZEB/miR-200 feedback loop showed significantly metastatic ability *in vivo* [93]. Future efforts are needed to dissect the functional consequences of breaking feedback loops among different axes of plasticity (such as the ones investigated here) in eventually delaying or preventing metastasis.

## Methods

### RAndom CIrcuit Perturbation (RACIPE)

RACIPE [43] is a tool allowing a parameter-agnostic computational simulation of regulatory networks based solely on the network topology. The network is composed of nodes representing various gene products and directed links representing inhibitory and activating links between them. For each node *i* with its expression level at a time *t* as p_*i*_, with incoming links from a set {A}of nodes, we have the Ordinary Differential Equation for the dynamics of p_*i*_ thus:

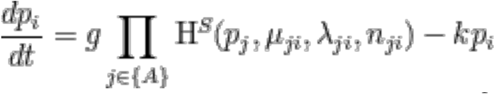

where *g* is the basal production rate, *k* is the degradation rate, *H*^*S*^ is the Shifted Hill-function with parameters *n, μ, λ, p* representing the effect of regulation by node *j* on *i*, altering its production rate. *λ* > 1 represents an activating link (H^*S+*^) and *λ* < 1represents an inhibitory link (H^*S*–^) with *λ* = 1 representing no regulation. *μ* is the threshold parameter that controls the level of *p* around which the non-linearity in *H*^*S*^ is seen. *n* is the Hill-coefficient representing the “extent of non-linearity” in the function by modulating the steepness of the response curve. See **Supplementary figure S9** for graphs of the Shifted Hill function for varied parameters.

### Parameter Sampling

Given a network topology, RACIPE generates an ensemble of random models each with a randomly sampled set of parameters for every link. The default range of random sampling has been estimated from BioNumbers [94], and here we choose the default option of uniform random sampling from the range. Instead of directly randomizing the basal production rate parameter *g*, RACIPE randomizes the maximal production rate *G*, following

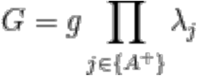

where {A^+^} is the set of all incoming activating links (production rate when all activating links are maximally active and all inhibitory links are inactive).

RACIPE allows perturbing the circuit by changing the expression levels of different nodes relative to the other unperturbed nodes. Overexpression and downregulation by *x* fold changes the sampling range for *G* by *x* times, hence, a 10-fold overexpression of a node will sample the *G* of the node from a range 10 times the range of the unperturbed (normal) node, while a 10-fold down-expression shrinks the range to 1/10th of its normal range. Details of the default ranges and the ranges used in our simulation are in **Supplementary section 1.1.2**.

### Simulation

We generate an ensemble of 10000 models with each run (analogous to a biological replicate, and referred elsewhere as such), and repeat this for 5 replicates to get the standard deviation for proportion (or frequency) calculations. RACIPE for each of the models simulates the network starting from multiple random initial positions until the network reaches a steady state, and prints as output the levels of each node at all identified steady-state solutions. The details of the run parameters used are in **Supplementary section 1.1.1 and 1.1.2**. The output of RACIPE for levels of all nodes are in log2 scale, and we perform a gene-wise normalization of the values before further analysis. For p_*i*_ representing the actual steady state levels of node *i*, the final values *z*_*i*_ are obtained after normalizing the log-scale node levels:

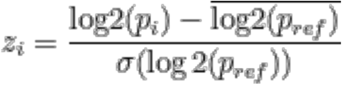

Here,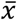 and *σ*(*x*)represents the mean-value and the standard deviation of the respective quantities. *p*_*ref*_ is taken as the corresponding node in the circuit with no over-expressed or down-expressed nodes (the “reference circuit”) allowing a uniform normalization for circuits across perturbations (see **Supplementary section 1.2.3**).

### Link Strength metrics

We define a link-strength metric similar to the one used earlier [95] to quantify the strength of each link independent of its activation at steady state. The link strength (*l*_*ij*_) is directly proportional to an activating *λ*^+^ or inversely proportional to an inhibitory *λ*^−^ and it is inversely proportional to the threshold μ. To normalize the threshold, we choose the normalization value *G/k*, the maximum possible node level at steady state.

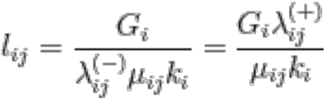

For any two links *l*_*ij*_ and *l*_*gh*_ (usually a part of a mutually-inhibitory feedback motif between two nodes), the asymmetry is defined as *log*_2_(*l*_*ij*_/*l*_*gh*_) representing the overall asymmetry in the effective strengths of the two links forming a mutually inhibitory feedback loop. When quantifying the coupling strength, the combined strength is defined as *log*_2_(*l*_*ij*_) + *log*_2_(*l*_*gh*_).

### Calculation of p1 and p2 values

For calculation of conditional probability of having a feature x conditional on having a feature y, we take the proportion of all steady state solutions having the feature y that have the feature x too. We use two such conditional probabilities in our results. p1 is the probability of lying in the stemness window conditional on the solution belonging to a particular cluster, and is calculated as the proportion of all solutions of the specific cluster lying in the stemness window. p2 is the probability of belonging to a particular cluster conditional on the solution lying in the stemness window, and is calculated as the proportion of all solutions in the stemness window belonging to a particular cluster.

### Clustering

For clustering, KMeans algorithm was used on the appropriately normalized data for only the four nodes (**miR-200, ZEB, LIN28**, and **let7**) with all solutions taken together with the same weight, without distinguishing between solutions belonging to systems with different number of stable states. KMeans finds an optimal splitting of the data into K clusters (where K is a hyperparameter that the algorithm takes as the input) by iteratively minimizing inertia, or the total within-cluster sum-of-squared distance from the cluster centres. K-Means results have a stochastic component since they depend upon the initialization of the cluster centres which are iteratively improved by the algorithm. To offset this, at each run, we randomize the initialization multiple times and select the best outcome in terms of minimized k-means inertia (See **Supplementary section 1.2.1** for further details of the algorithm parameters). In order to select a cluster number, we run the K-Means algorithm with different values for K and at each value, compute multiple cluster-quality metrics, all of which attempt to quantify the optimality of the clusters obtained at each value for K, and we then plot these metrics across different values of K. A sharp change in the slope, as well as an overall higher value for the average Silhouette widths indicate an appropriate partitioning [96]. For our data, the peak and the sudden change in slope [97] were almost unambiguously in the favour of K=4 in all the metrics that we used (Average Silhouette widths, Calinski-Harabasz index [98], Davies-Bouldin index [99], K-Means inertia) (see **Supplementary section 1.2.2** and **Fig S2**). Other relatively subjective supportive evidence can be gained from the visual evaluation of the PCA plot and observing the dendrogram computed using agglomerative hierarchical clustering using Ward’s minimum variance criterion [100]. The clustering algorithms and the cluster-quality metrics were from scikit-learn version 0.32 (see **Supplementary section 1.1.3** for more details on the software versions and the platform). Each replicate was separately clustered and normalized (see **Supplementary section 1.2.3** for details on normalization).

### Significance Testing

To compute the statistical significance for all comparisons, we have used two strategies. For comparisons within the same RACIPE replicate (where n is large enough), we use the two-sided non-parametric Mann-Whitney U Test [101] (also known as the Wilcoxon rank-sum test). For cross-replicate comparisons (i.e. proportions), we use the Welch t-test [102] allowing unequal variances though with the assumption of normality of the compared groups. Mann-Whitney U test is not suitable for small samples as it discards all information about the size of the differences (and as a result, caps the highest significance that can be achieved based on n) and also the assumption of normality of the proportions across replicates is a relatively reasonable assumption since it is stochastic differences which primarily cause the inter-replicate values to differ. Furthermore, we employ the Holm-Bonferroni correction to adjust the p-values accounting for multiple comparisons to control a Family-Wise Error Rate (FWER) in the overall rejection of the null hypotheses (see **Supplementary section 2.5**). Due to the large number of steady states per RACIPE replicate, for some comparisons within the same replicate, very small differences of means (or medians) gives a statistically significant p-value but not necessarily biologically relevant. Thus, to ascribe possible biological meaning to such comparisons, for cases of irrelevantly small “magnitude” of this difference (i.e. the effect size), we provide the ratio of the means/medians and the absolute difference of these quantities.

## Acknowledgements

This work was supported by Ramanujan Fellowship awarded to MKJ by Science and Engineering Research Board (SERB), Department of Science and Technology (DST), Government of India (SB/S2/RJN-049/2018).

## Author contributions

MKJ conceived and supervised research; SP performed research; SP and SS analysed data; all authors contributed to writing and editing of the manuscript.

## Code availability

All relevant codes are available athttps://github.com/Stochastic13/emt-stemness

## Supplementary Figures

**Figure S1:**
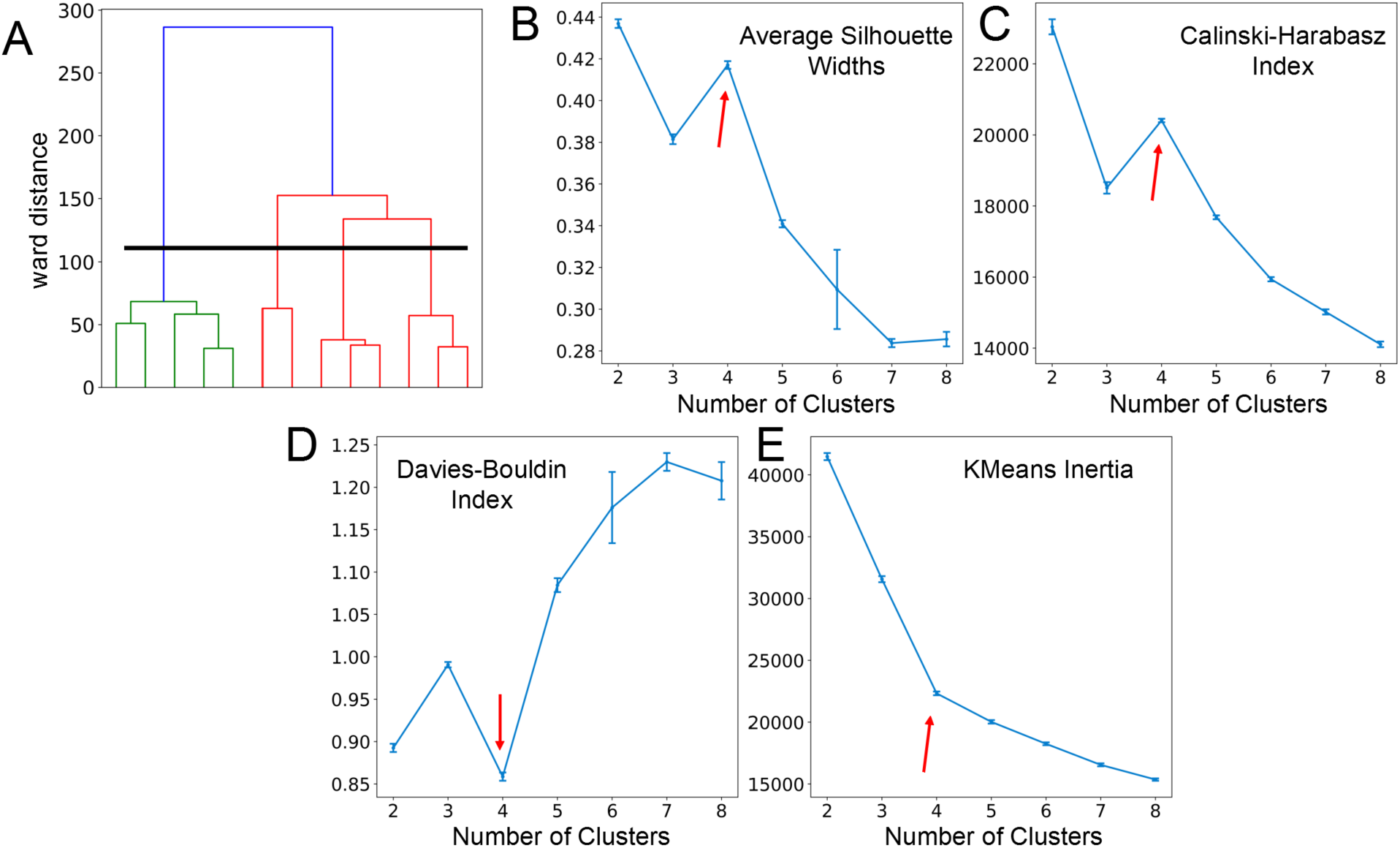
Optimal cluster number by various cluster quality. **A)** Dendrogram of a single replicate of the coupled EMP-stemness circuit computed using the ward-algorithm under agglomerative hierarchical clustering. The black horizontal line represents segregation of dendrogram corresponding to a 4-cluster solution. **B)** Average Silhouette widths for the coupled EMP-stemness circuit at different numbers of clusters with a distinct peak (and a marked change in the slope) at cluster number = 4. The larger the average Silhouette width, the more optimal is the clustering. **C)** Calinski-Harabasz index for the coupled EMP-stemness circuit at different numbers of clusters with a distinct peak (and a marked change in the slope) at cluster number = 4. The larger the index, the more optimal is the clustering. **D)** Davies-Bouldin index for the coupled EMP-stemness circuit at different numbers of clusters with a distinct dip (and a marked change in the slope) at cluster number = 4. The smaller the index, the more optimal is the clustering. **E)** K-means inertia (the objective of the K-means algorithm is to minimize this quantity iteratively) for the base circuit at different numbers of clusters with a distinct “elbow” (and a marked change in the slope) at cluster number = 4. All the values are averaged across n=5 RACIPE replicates and the error bars represent the across-replicate standard deviations.

**Figure S2:**
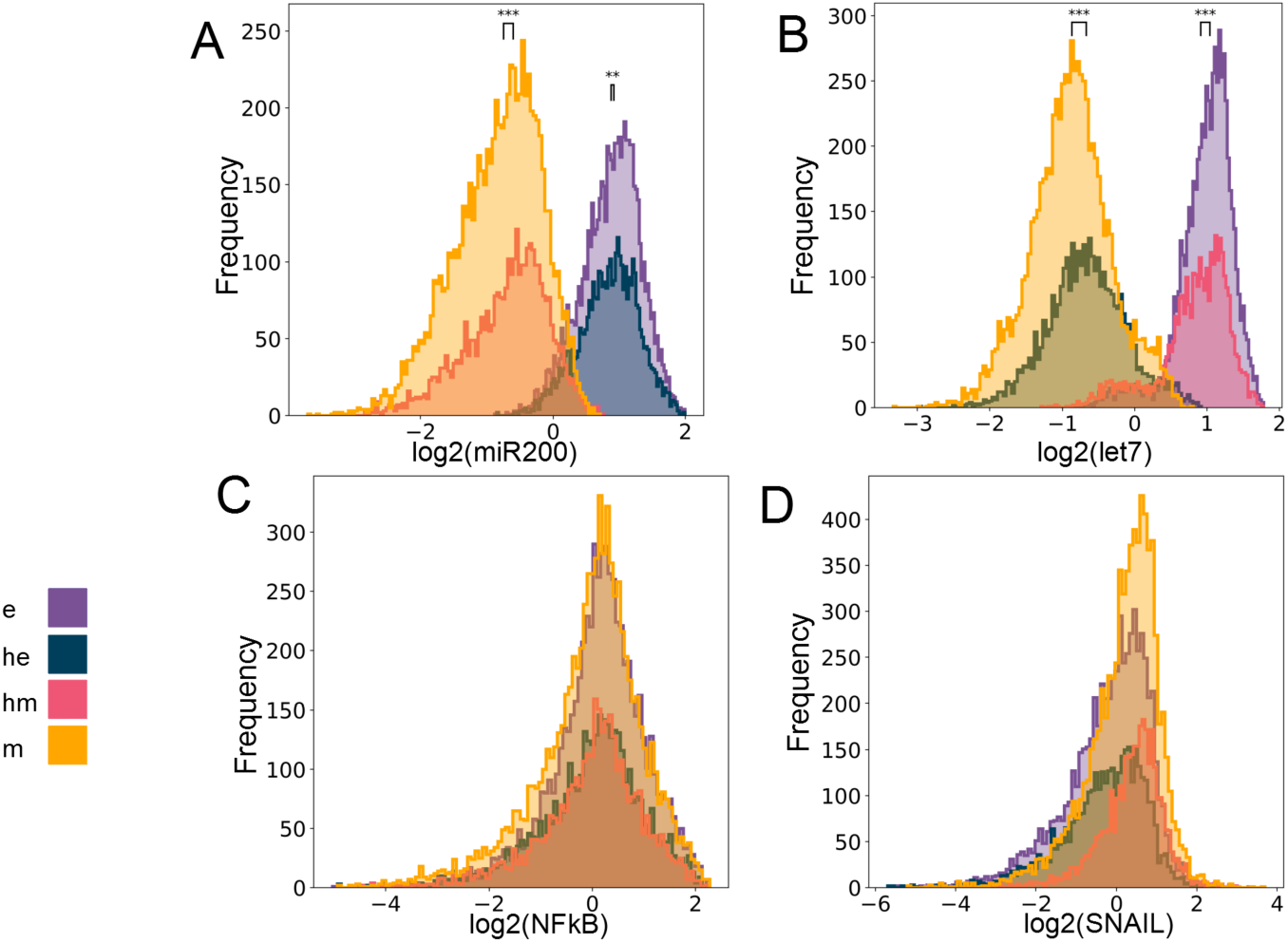
Four E/M phenotypes enabled by the Base circuit. **A-D)** Expression histograms showing miR-200, let7, NF-kB and SNAIL levels respectively in the 4 clusters – e, he, hm and m. Significance bars for statistical testing (Mann-Whitney U test) added in B and C. (legend for the p-values +: p > 0.01, *: 0.01 > p > 0.001, **: 0.001 > p > 0.0001, ***: p < 0.0001; for effect sizes refer to **Table S7** in **Supplementary Section 2.5**)

**Figure S3:**
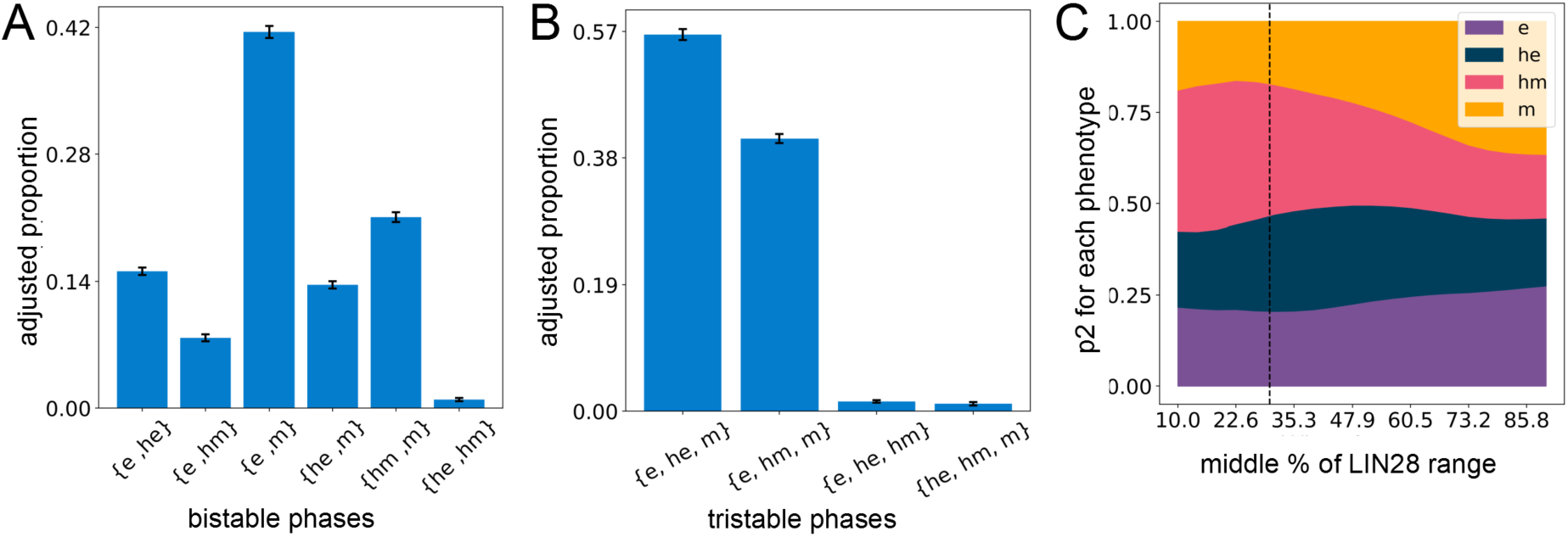
Stemness Window and the bistable/tristable phase distributions of the coupled EMP-stemness circuit. **A)** Proportion of all bistable parameter sets in the coupled EMP-stemness (base) circuit belonging to different bistable phases, after removing the parameter sets with both solutions being assigned the same phenotype (see **Supplementary section 2.3**). **B)** Same as A) but for tristable systems, after removing the systems with 2 or more solutions being assigned the same phenotype. **C)** Sensitivity analysis by considering multiple sizes of the stemness windows centered at the median of the LIN28 distribution (averaged across n=5 RACIPE replicates of the base circuit). The proportion of all high-stemness solutions belonging to each of the phenotypes (p2) is then plotted. The black vertical line corresponds to the 30% value taken as the stemness window and marks the approximate upper limit of the range with a relatively small change in the phenotype-distribution upon altering the stemness window (A, B: The values are averaged across n=5 RACIPE replicates and the error bars represent the across-replicate standard deviations.)

**Figure S4:**
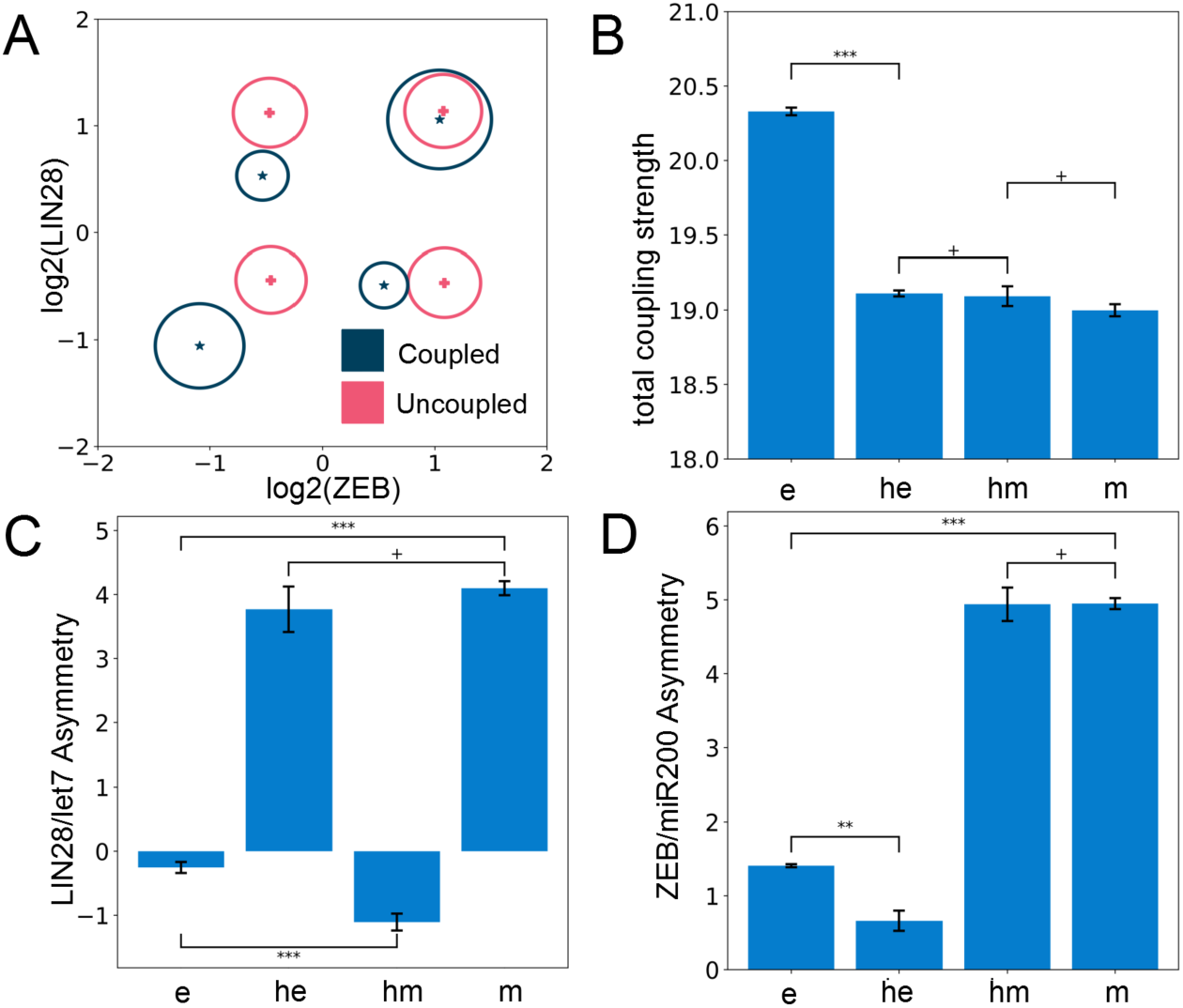
Link Strength analysis. **A)** The cluster centers (median levels of ZEB and LIN28 s) in the ZEB-LIN28 plane for the coupled and the uncoupled EMP-stemness circuits. The radius of the circle around the center is proportional to the proportion of all solutions falling into the cluster. Both the uncoupled and the base circuit have been normalized with the coupled EMP-stemness circuit (see **Supplementary Section 2.2**). **B)** The total coupling strength (see **methods**) plotted for different phenotypes in the base circuit including only the parameter sets resulting in a monostable solution. **C)** Mean symmetry ratio (see **methods**) of the LIN28/let-7 double negative feedback loop plotted for different phenotypes in the base circuit including only the parameter sets resulting in a monostable solution. A higher ratio represents a stronger suppression of let7 by LIN28 in comparison to the reverse inhibition. **D)** Same as C) but for ZEB/miR-200 feedback loop. A higher ratio represents a stronger suppression of miR200 by ZEB in comparison to the reverse inhibition. (B-D: The values are averaged across replicates and the error bars represent the across-replicate standard deviations. Welch’s t-test is used significance. Legend for the p-values +: p > 0.01, *: 0.01 > p > 0.001, **: 0.001 > p > 0.0001, ***: p < 0.0001; for effect sizes refer to **Table S7** in **Supplementary Section 2.5**)

**Figure S5:**
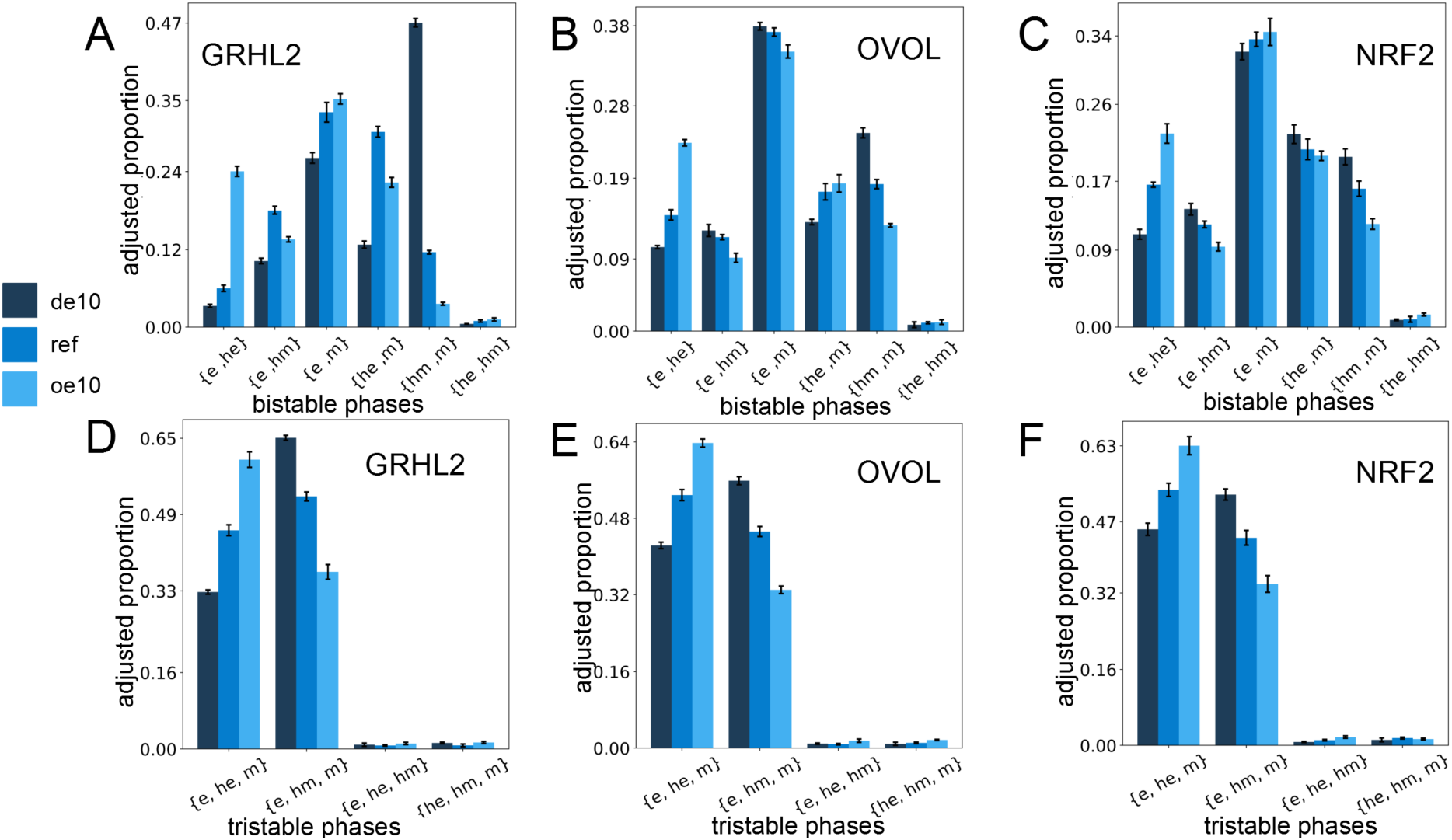
Distribution of bistable and tristable systems upon incorporating PSFs with the coupled EMP-stemness circuit. **A-C)**: Frequency of different possible combinations of co-existing bistable states (i.e. phases) with varied expression of the PSFs for different circuits. Y-axis represent the proportion of all bistable systems belonging to different bistable phases after removing the systems with both solutions being assigned the same phenotype (See **Supplementary section 2.3**). **D-F)** Same as A-C but for tristable solutions after removing the systems with 2 or more solutions being assigned the same phenotype (de10: 10-fold down-expression, ref: reference unperturbed circuit, oe10: 10-fold overexpression). The values shown here are averaged across n=5 RACIPE replicates and the error bars represent the across-replicate standard deviations.

**Figure S6:**
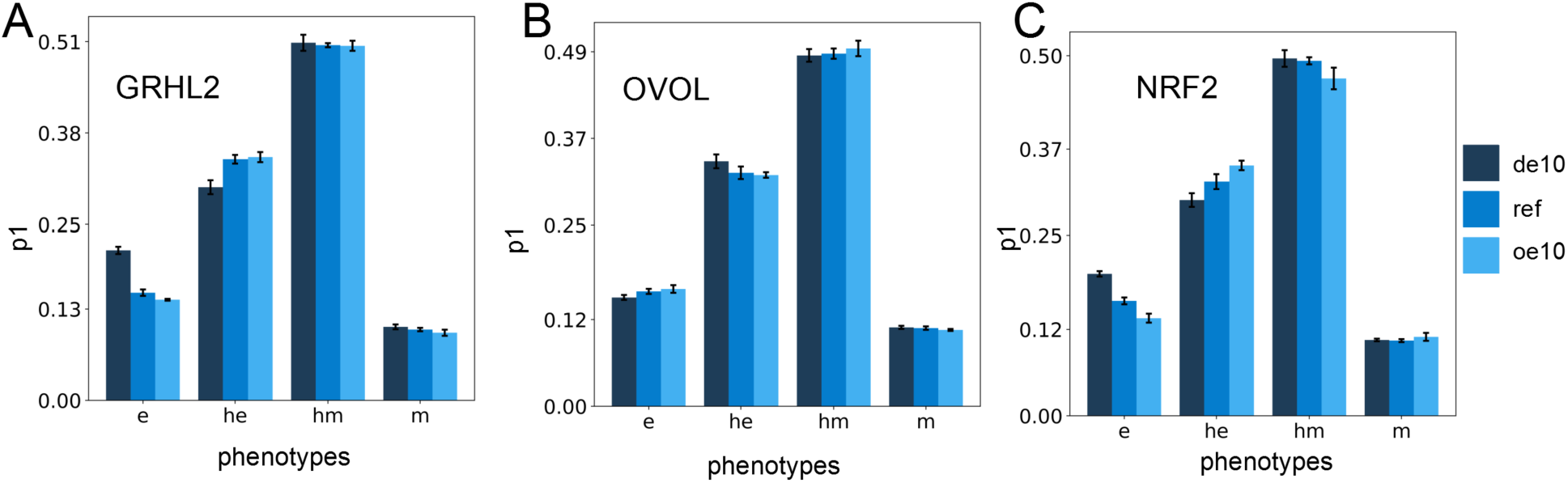
Stemness probability (p1) changes with PSF expression. **A-C)** Changes in the p1 probability for different PSF circuits at different expression levels of PSFs. See **Table S8** in **Supplementary Section 2.5** for the statistical testing for differences. (de10: 10-fold downexpression, ref: reference unperturbed circuit, oe10: 10-fold overexpression) (The values are averaged across n=5 RACIPE replicates and the error bars represent the across-replicate standard deviations)

**Figure S7:**
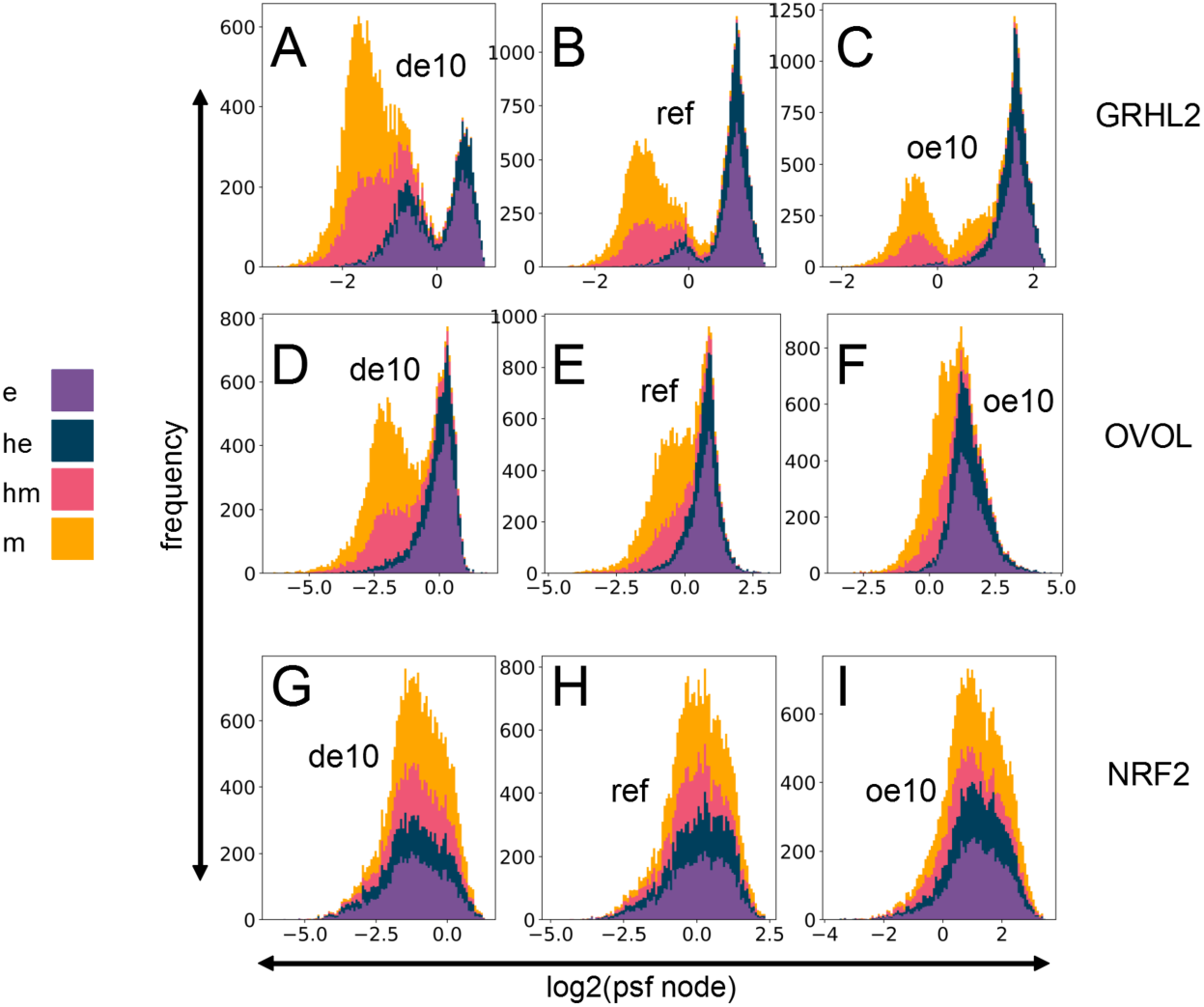
PSF expression profiles at different expression levels. **(A-C)** GRHL2 stacked histograms for a single RACIPE replicate (colored according to the phenotypes as shown in the legend) for different levels of expression of GRHL2 node. **(D-F)** Same as A-C but for OVOL. **(G-I)** Same as A-C but for NRF2 (de10: 10-fold downexpression, ref: reference unperturbed circuit, oe10: 10-fold overexpression)

**Figure S8:**
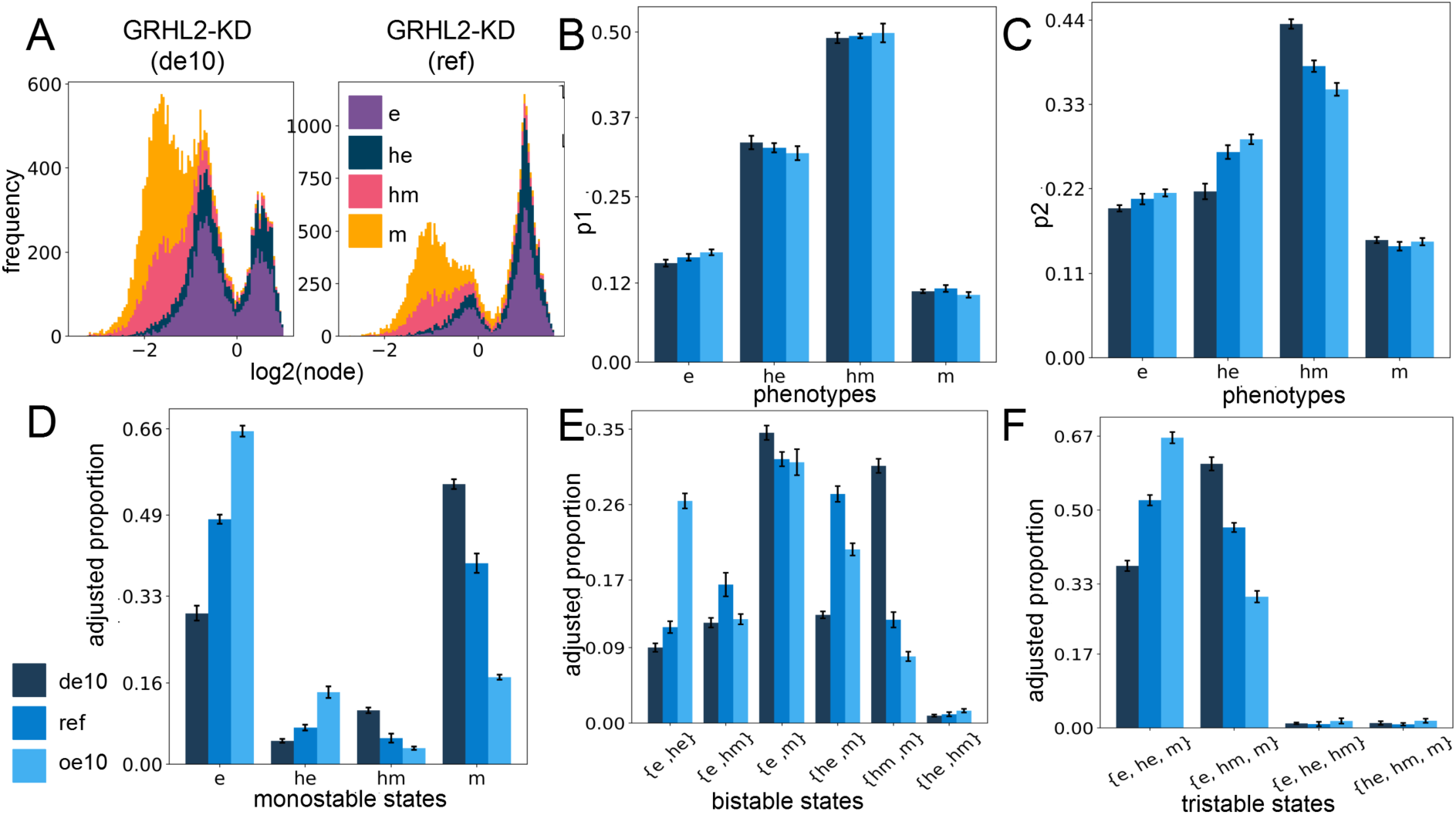
GRHL2-KD circuit results for the investigated quantities. **A)** Stacked histograms for GRHL2 in one replicate of GRHL2-KD circuit at 10-fold downexpression (left panel) and the unperturbed circuit (right panel) with different phenotypes colored separately. **B)** Same as Fig S6A-C but for GRHL2-KD circuit, with different levels of expression of the GRHL2 node. **C)** Same as Fig 4C-E but for GRHL2-KD circuit, with different levels of expression of the GRHL2 node. **D)** Proportion of all states from monostable parameter sets belonging to different phenotypes in the GRHL2-KD circuit at different levels of expression of the GRHL2 node. **E**,**F)** Same as Fig S5 but for GRHL2-KD circuit, with different levels of expression of the GRHL2 node (B-F: The values are averaged across replicates and the error bars represent the across-replicate standard deviations; for B-D see **Table S8** in **Supplementary section 2.5** for the statistical testing for differences).

**Figure S9:**
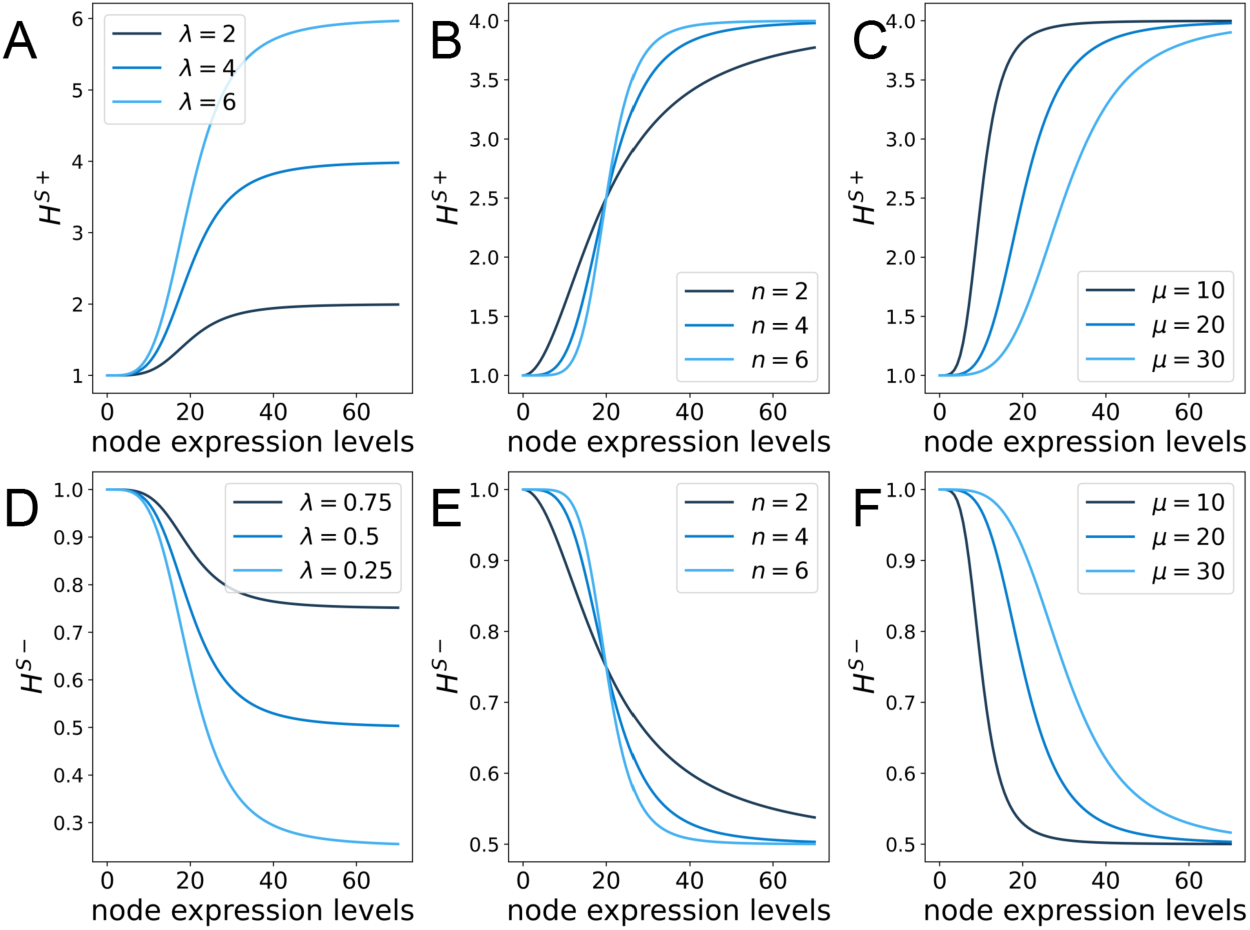
Shifted Hill Function across different parameters. **(A-C)** Shifted Hill Function for an activating link. A: μ=20, n=4 and varying λ, B: μ=20, varying n and λ=4, C: varying μ, n=4 and λ=4. **(D-F)** Shifted Hill Function for an inhibiting link. D: μ=20, n=4 and varying λ, E: μ=20, varying n and λ=0.5, F: varying μ, n=4 and λ=0.5

