## Supplementary Information for "Hybrid E/M phenotype(s) and stemness: a mechanistic connection embedded in network topology"

### 1 Methods

#### 1.1 RACIPE

##### 1.1.1 Simulation details

The final run options include 10000 random parameter sets and 200 initial values per parameter set with Euler's method for integration. The threshold (minimum euclidean distance between two solutions to consider them separate), step-size (for integration) and maximum number of iterations (to reach steady state) were kept at their default values (see **Table S1**). In order to select the simulation options, runs with different parameters were performed for the base circuit, and the stability of the results compared. The results were stable even with high-accuracy parameters (Runge-Kutta method for integration, lower step size and a lower threshold), hence the default options were chosen to minimize computation time.

##### 1.1.2 Parameter Ranges

The default parameter ranges[1] and the parameter ranges used in the reference unperturbed circuit are given in the **Table 1**. The $G$ parameter is appropriately scaled in the down-expression and the over-expression circuits with the remaining parameters (and simulation options) kept the same.

| parameter | default range | simulation range |
|---|---|---|
| $G$ | 1-100 | 1-500 |
| $k$ | 0.1-1 | 0.1-1 |
| $\lambda^{(+)}$ | 1-100 | 1-100 |
| $1/\lambda^{(-)}$ | 1-100 | 1-100 |
| $\mu$ | half-functional | half-functional |
| $n$ | 1-6 | 1-6 |
| number of models | 100 | 10000 |
| threshold | 1 | 1 |
| stepsize | 0.01 | 0.01 |
| maxiters | 20 | 20 |
| num-ode | 100 | 200 |
| num-stability | 10 | 15 |
| dist | uniform | uniform |
| num-find-T | 10000 | 10000 |

**Table S1:** Parameters ranges and the simulation options

The range for $G$ has been changed from the default upper limit of 100 to 500, because, RACIPE auto-amplifies the range of max-production (G) for certain nodes based on the number of incoming links in order to get an appropriate value for the half-functional threshold. This may lead to the max-production ranges of some nodes to be relatively different, which might make inference difficult. Setting the reference range to the higher value removes this auto-amplification allowing a uniform range for all nodes in the reference circuit without disturbing any of our other assumptions since we perform a gene-wise normalization before analysis. $x$-fold over-expression (or down-expression) expands (or shrinks) this new range $x$ times. The increase in the num-stability parameter is only to obtain a more detailed division of the number of stable states, without much relevance for the rest of the results.

##### 1.1.3 Platform Details

All simulations were performed on an Acer laptop with Intel core i7 (8th Generation) with Windows 10. RACIPE was compiled using **GNU make** (v4.3) and **gcc** (v9.3.0) on **cygwin** (v3.1.6). The versions of python libraries used were as follows: **python interpreter** v3.6.8 (the main python interpreter), **numpy** v1.18.4 (basic manipulations), **pandas** v1.0.3 (basic manipulations), **scipy** v1.4.1 (primarily for statistical tests and dendrogram), **sci-kit learn** v0.23.1 (for clustering algorithms, PCA, cluster quality metrics), **matplotlib** v3.2.1 (primary plotting library), **seaborn** v0.11.0 (for heatmap), **statsmodels** v0.11.1 (for multiple correction in statistical testing).

#### 1.2 Clustering

##### 1.2.1 algorithm details

KMeans implementation of scikit-learn allows an "intelligent" initialization of the initial cluster centers by choosing one of the data points randomly as one of the cluster centers and then initializing the rest by trying to maximize the separation between them, and this may lead to better results. We use this strategy in our initialization (called *kmeans++*[2] in literature and the algorithm implementation). For each cluster number, 50 initial positions are initiated and the solution with a minimum inertia is chosen as the final output. The algorithm attempts a maximum of 500 iterations to reach convergence with a tolerance of $10^{-5}$ (i.e. solution is considered to have converged if the improvement on next iteration is below the tolerance).

##### 1.2.2 Quality Metrics

Since the task of choosing an appropriate cluster number is a hard task, often involving a great deal of subjectivity[3, 4], for our purposes, we employ multiple different lines of relatively objective criteria to select a cluster number. Primary among these are what we refer here as "cluster quality metrics", or metrics commenting on the optimality of the clustering partition. Of them, silhouette widths[3] are the easiest to interpret in terms of the underlying mathematical logic, while the other criteria used have a relatively more complicated underlying basis. Silhouette width for a single sample is defined as follows(where $a$ is the mean intra-cluster distance for a sample, and $b$ is the mean nearest-cluster distance for a sample):

$$\frac{b-a}{\max(a,b)}$$

This width can vary between -1 and 1, with negative values indicating a wrong cluster assignment (nearest cluster being closer to the sample than its own assigned cluster), while a value close to 0 represents an "artificial" division between the self-cluster and the nearest cluster according to the sample. We take an all-sample average width as our criterion. However, for all of the criteria used,

a sharp change in value is considered a good measure of optimality of the obtained clusters as well as the direct maximization (silhouette widths, Calinski-Harabasz index[5]) or minimization (Davies-Bouldin index[6]) of the metric. In KMeans inertia, the elbow method (similar to a sharp change in slope) is most relevant since unlike the other metrics which penalize closely spaced clusters (i.e. artificial division), inertia monotonically decreases in most cases with increasing cluster number. For our cases, all of these metrics and all the forms of evaluating them (change in slope vs maximization/minimization) universally report a consistent value of 4 clusters in all the circuits. **Fig S2** shows a combination of all the 4 indices as used for the base circuit.

##### 1.2.3 Normalization

We perform a run-wise gene-specific normalization for each circuit using the unperturbed circuit as reference. In the base circuit, where there is no perturbation, each run is separately normalized. For all the PSF circuits, each run number x of the down-expressed and the over-expressed circuits are normalized with reference to run number x of the unperturbed circuit. The rationale behind separate normalization of the base and the PSF circuits is that in RACIPE, every node's max-production range depends on the incoming activating links, which may differ between circuits, hence rendering them not directly comparable. Instead, over-expressing and down-expressing a node in the same topology makes inferences much more relevant and attributable to the effect of that particular node.

### 2 Results

#### 2.1 Stability of the RACIPE replicates

Illustrations supporting qualitative features of the node level distributions across clusters (histograms, boxplots, PCA) are considered for a single replicate in the main article. While the cross-replicate standard deviations in case of calculated proportions suffice for establishing the stability of the concerned proportions across the replicates, we need some other quantification to understand the stability of these cluster distributions. These were also found to be very stable across replicates (**Table S2**) as indicated by a stable median and inter-quartile range. These were chosen to be the statistics for central tendency and dispersion respectively since the distributions may not be unimodal or symmetrical making means and standard deviations difficult to interpret.

| Replicate (node) | e | he | hm | m |
|---|---|---|---|---|
| 1 (ZEB) | -1.09 (0.69) | -0.531 (0.559) | 0.547 (0.48) | 1.04 (0.383) |
| 2 (ZEB) | -1.1 (0.678) | -0.55 (0.555) | 0.527 (0.52) | 1.04 (0.377) |
| 3 (ZEB) | -1.1 (0.68) | -0.545 (0.558) | 0.534 (0.524) | 1.04 (0.364) |
| 4 (ZEB) | -1.09 (0.679) | -0.528 (0.582) | 0.543 (0.518) | 1.04 (0.38) |
| 5 (ZEB) | -1.1 (0.684) | -0.541 (0.577) | 0.555 (0.496) | 1.05 (0.377) |
| 1 (LIN28) | -1.06 (0.675) | 0.532 (1.04) | -0.49 (0.604) | 1.06 (0.491) |
| 2 (LIN28) | -1.07 (0.675) | 0.509 (1.09) | -0.503 (0.615) | 1.05 (0.488) |
| 3 (LIN28) | -1.07 (0.668) | 0.514 (1.06) | -0.485 (0.631) | 1.07 (0.481) |
| 4 (LIN28) | -1.07 (0.664) | 0.523 (1.05) | -0.48 (0.634) | 1.06 (0.483) |
| 5 (LIN28) | -1.07 (0.663) | 0.516 (1.05) | -0.471 (0.641) | 1.06 (0.495) |

**Table S2:** Median (iqr) for ZEB and LIN28 in different replicates of the base circuit rounded off to 3 significant figures

#### 2.2 Uncoupled Circuit

In order to simulate the uncoupled circuit, we ran two separate RACIPE runs, one for the E/M module (ZEB, miR-200, SNAIL) and another for the stemness module (LIN28, let-7, NFkB). Each of the separate runs give a 2-cluster solution (**Table S3**. RACIPE being a coarse-grained view of the dynamics allows robust interpretation at the cost of finer structure not being easily appreciable. Hence, the information of the hybrids may be lost in the overlap of the two clusters or the specific tristable systems in the ensemble of the obtained solutions. It may also be explained by the absence of the complexity of miR regulation which is replaced here by a simple shifted-Hill function based regulation without any separate mRNA nodes). After the cluster assignment for each of the circuits individually, since they are independent circuits, any of the parameter sets from one circuit can be paired with any of the parameter sets of the other circuit with equal likelihood. Instead of the parameter-set pairing, we randomly pair up the obtained solutions choosing a simpler procedure. We then naturally get 4 clusters from the combined circuit (2 from each randomly paired with the 2 from the other circuit).

| number of clusters | E/M module | stemness module |
|---|---|---|
| 2 | 0.63 | 0.61 |
| 3 | 0.52 | 0.53 |
| 4 | 0.41 | 0.52 |
| 5 | 0.40 | 0.39 |
| 6 | 0.37 | 0.38 |
| 7 | 0.36 | 0.37 |
| 8 | 0.37 | 0.37 |

**Table S3:** Average Silhouette widths for the Uncoupled E/M and Stemness circuits

#### 2.3 Multistable phases

In order to get the proportion of x-stable solutions (eg. bistable solutions) belonging to a particular x-stable phase (eg. {e, m}), we filtered out all x-stable parameter sets (giving x-stable systems) with more than one solution in the same cluster. For the base circuit, this filtering out does not have any significant impact on the inference, but it becomes more relevant while comparing the results across different expression levels of the PSFs in their respective circuit. This group of the filtered-out systems does not change in proportion significantly across expressions in the PSF circuits, hence allowing interpretations of the observed trends without confounding (**Table S4**)

| **circuit** | **bi** | **tri-2** | **tri-3** |
|---|---|---|---|
| base | 0.13 (0.0051) | 0.39 (0.011) | 0.0057 (0.0015) |
| GRHL2 (de10) | 0.16 (0.0044) | 0.47 (0.016) | 0.0095 (0.0017) |
| GRHL2 (ref) | 0.13 (0.0057) | 0.38 (0.0074) | 0.0061 (0.0013) |
| GRHL2 (oe10) | 0.13 (0.0081) | 0.41 (0.0082) | 0.011 (0.0016) |
| OVOL (de10) | 0.13 (0.012) | 0.35 (0.0037) | 0.0073 (0.0011) |
| OVOL (ref) | 0.13 (0.0049) | 0.36 (0.013) | 0.0065 (0.0017) |
| OVOL (oe10) | 0.13 (0.007) | 0.38 (0.014) | 0.0077 (0.0019) |
| NRF2 (de10) | 0.14 (0.0047) | 0.35 (0.015) | 0.0072 (0.0017) |
| NRF2 (ref) | 0.15 (0.0059) | 0.4 (0.015) | 0.01 (0.0022) |
| NRF2 (oe10) | 0.15 (0.0088) | 0.4 (0.0044) | 0.009 (0.0015) |
| GRHL2-KD (de10) | 0.2 (0.0066) | 0.47 (0.015) | 0.017 (0.0026) |
| GRHL2-KD (ref) | 0.25 (0.0071) | 0.5 (0.0068) | 0.021 (0.0025) |
| GRHL2-KD (oe10) | 0.21 (0.0048) | 0.48 (0.0092) | 0.025 (0.0035) |

**Table S4:** Proportion (averaged across all replicates) of filtered x-stable parameter sets based on $> 1$ solutions of the set being assigned the same phenotype (standard deviations in the parentheses). "bi": proportion of bistable sets with 2 solutions of the same phenotype, "tri-2": tristable sets with 2 solutions of the same phenotype and "tri-3": tristable sets with 3 solutions of the same phenotype.

#### 2.4 Overall Solution Distribution

The distributions of all solutions (irrespective of the number of states of the corresponding parameter set) across the phenotypes (**Table S5**) form an underlying basis to interpret the p1 and p2 plots in the main article. However, they are a relatively poorer proxy measure for the "probabilities" of existence of these phenotypes, since states with very small attractor basins (i.e. the proportion of all random initializations converging to this state) compared to other states of the same parameter set are considered together with the same weight. Alternatively, considering only monostable solutions removes this bias as done in the main article.

| circuit | e | he | hm | m |
|---|---|---|---|---|
| base | 0.3 (0.0036) | 0.18 (0.0013) | 0.17 (0.0048) | 0.35 (0.0034) |
| GRHL2 (de10) | 0.24 (0.0018) | 0.11 (0.0025) | 0.26 (0.0014) | 0.39 (0.0031) |
| GRHL2 (ref) | 0.3 (0.0025) | 0.19 (0.0027) | 0.19 (0.0022) | 0.32 (0.0012) |
| GRHL2 (oe10) | 0.31 (0.0016) | 0.22 (0.0019) | 0.14 (0.0029) | 0.33 (0.0022) |
| OVOL (de10) | 0.29 (0.0032) | 0.15 (0.0018) | 0.21 (0.0022) | 0.36 (0.0035) |
| OVOL (ref) | 0.3 (0.0019) | 0.18 (0.002) | 0.18 (0.00038) | 0.34 (0.0019) |
| OVOL (oe10) | 0.32 (0.0021) | 0.21 (0.0025) | 0.15 (0.0028) | 0.31 (0.0026) |
| NRF2 (de10) | 0.27 (0.003) | 0.18 (0.0029) | 0.21 (0.0012) | 0.34 (0.0026) |
| NRF2 (ref) | 0.28 (0.0038) | 0.21 (0.0039) | 0.19 (0.0029) | 0.32 (0.0041) |
| NRF2 (oe10) | 0.3 (0.0016) | 0.22 (0.0041) | 0.17 (0.0054) | 0.31 (0.0024) |
| GRHL2-KD (de10) | 0.3 (0.0018) | 0.15 (0.0032) | 0.21 (0.0021) | 0.34 (0.0021) |
| GRHL2-KD (ref) | 0.31 (0.0023) | 0.2 (0.0022) | 0.18 (0.0031) | 0.31 (0.0025) |
| GRHL2-KD (oe10) | 0.29 (0.0015) | 0.21 (0.0021) | 0.16 (0.0012) | 0.34 (0.0021) |

**Table S5:** Proportion (averaged across all replicates) of all solutions irrespective of the number of stable states of the corresponding parameter set belonging to different phenotypes (with standard deviations in parenthesis)

#### 2.5 Statistical Testing

While the number of hypotheses constituting the different "hypotheses-families" for multiple-testing correction is quite small in our study, we are still performing the Holm-Bonferroni[**?**] correction to control a family-wise error rate (fwer). Choosing these "families" is often subjective. Here we club all the testing done between two same data groups into a single "family" in accordance with the common practice, with the theoretical independence of all hypotheses being a practically acceptable assumption (as long as there are no major dependencies between the hypotheses of a single group). In **Table S6** and **Table S7** color coding represents a semi objective assessment for a relevant effect size. For example, in **Table S7** a relevant effect size includes a ratio $> 1.2$ or $< 0.8$ and an absolute difference $\geq 0.5$. The chosen $\alpha$ for significance is 0.01 in the following tests. An additional measure of the effect size (the absolute difference of medians) is considered here since the groups can be centered around zero and the ratio of medians with a differing sign overlooks the actual magnitude of the difference. Furthermore, for medians near zero, a small absolute change may result in a large ratio. In **Table S9**, a Welch t-test is done between the PSF circuit with 10-fold down-expression (de10) and the circuit with 10-fold overexpression (oe10); proportions calculated for the five replicates of each of these perturbed circuits forming one comparison group. This allows performing a statistical test for the enrichment of different quantities due to the increased expression of the PSF node.

| parameter | group 1 | group 2 | p-value | $m1/m2$ | $\|m1 - m2\|$ |
|---|---|---|---|---|---|
| monostable[3] | e | m | + | 1.0 | 0.017 |
| monostable[4] | he | hm | $*$ | 1.2 | 0.016 |
| p1[1] | e | he | $***$ | 0.46 | 0.18 |
| p1[3] | e | m | $***$ | 1.4 | 0.045 |
| p1[4] | he | hm | $***$ | 0.69 | 0.15 |
| p1[6] | hm | m | $***$ | 4.4 | 0.38 |
| p2[1] | e | he | $***$ | 0.79 | 0.055 |
| p2[3] | e | m | $**$ | 1.2 | 0.034 |
| p2[4] | he | hm | $***$ | 0.72 | 0.1 |
| p2[6] | hm | m | $***$ | 2.1 | 0.19 |
| ZEB-miR200 asymmetry[1] | e | he | $**$ | 2.1 | 0.75 |
| ZEB-miR200 asymmetry[3] | e | m | $***$ | 0.28 | 3.5 |
| ZEB-miR200 asymmetry[6] | hm | m | + | 1.0 | 0.0096 |
| LIN28-let7 asymmetry[2] | e | hm | $***$ | 0.23 | 0.85 |
| LIN28-let7 asymmetry[3] | e | m | $***$ | -0.062 | 4.4 |
| LIN28-let7 asymmetry[5] | he | m | + | 0.92 | 0.33 |
| Total coupling strength[1] | e | he | $***$ | 1.1 | 1.2 |
| Total coupling strength[4] | he | hm | + | 1.0 | 0.018 |
| Total coupling strength[6] | hm | m | + | 1.0 | 0.095 |

**Table S6:** Statistical Testing results for differences of various calculated proportions in the base circuit considering all the replicates (i.e. the proportion calculated for the 5 replicates forms one comparison group). $m1$, $m2$ are the means for group 1 and 2 respectively. (Welch t-test; +: $p > 0.01$, $*$: $0.01 > p > 0.001$, $**$: $0.001 > p > 0.0001$, $***$: $p < 0.0001$, green represents a significant difference and a relevant effect size, yellow indicates a significant difference without a relevant effect size and red indicates a non-significant result) The groups of hypotheses adjusted for multiple correction together are marked by unique subscript numbers.

| parameter | group 1 | group 2 | p-value | $m1/m2$ | $\|m1-m2\|$ |
|---|---|---|---|---|---|
| $ZEB^1$ | e | he | $***$ | 2.1 | 0.56 |
| $ZEB^2$ | e | hm | $***$ | -2.0 | 1.6 |
| $ZEB^3$ | e | m | $***$ | -1.0 | 2.1 |
| $ZEB^4$ | he | hm | $***$ | -0.97 | 1.1 |
| $ZEB^5$ | he | m | $***$ | -0.51 | 1.6 |
| $ZEB^6$ | hm | m | $***$ | 0.52 | 0.5 |
| $miR200^1$ | e | he | $**$ | 1.0 | 0.042 |
| $miR200^2$ | e | hm | $***$ | -1.5 | 1.5 |
| $miR200^3$ | e | m | $***$ | -1.2 | 1.7 |
| $miR200^4$ | he | hm | $***$ | -1.4 | 1.5 |
| $miR200^5$ | he | m | $***$ | -1.2 | 1.6 |
| $miR200^6$ | hm | m | $***$ | 0.81 | 0.15 |
| $LIN28^1$ | e | he | $***$ | -2.0 | 1.6 |
| $LIN28^2$ | e | hm | $***$ | 2.2 | 0.57 |
| $LIN28^3$ | e | m | $***$ | -1.0 | 2.1 |
| $LIN28^4$ | he | hm | $***$ | -1.1 | 1.0 |
| $LIN28^5$ | he | m | $***$ | 0.5 | 0.53 |
| $LIN28^6$ | hm | m | $***$ | -0.46 | 1.5 |
| $let7^1$ | e | he | $***$ | -1.5 | 1.7 |
| $let7^2$ | e | hm | $***$ | 1.1 | 0.13 |
| $let7^3$ | e | m | $***$ | -1.2 | 1.9 |
| $let7^4$ | he | hm | $***$ | -0.73 | 1.6 |
| $let7^5$ | he | m | $***$ | 0.77 | 0.2 |
| $let7^6$ | hm | m | $***$ | -1.0 | 1.8 |
| $SNAIL^1$ | e | he | $***$ | 0.087 | 0.15 |
| $SNAIL^2$ | e | hm | $***$ | -0.029 | 0.5 |
| $SNAIL^3$ | e | m | $***$ | -0.042 | 0.35 |
| $SNAIL^4$ | he | hm | $***$ | -0.33 | 0.65 |
| $SNAIL^5$ | he | m | $***$ | -0.49 | 0.49 |
| $SNAIL^6$ | hm | m | $***$ | 1.5 | 0.15 |
| $NFkB^1$ | e | he | $***$ | 2.5 | 0.12 |
| $NFkB^2$ | e | hm | $***$ | 2.0 | 0.097 |
| $NFkB^3$ | e | m | $***$ | 2.3 | 0.11 |
| $NFkB^4$ | he | hm | + | 0.81 | 0.019 |
| $NFkB^5$ | he | m | + | 0.93 | 0.0061 |
| $NFkB^6$ | hm | m | + | 1.1 | 0.012 |

**Table S7:** Statistical Testing results for differences of the node distributions across the clusters for a single replicate of the base circuit. $m1$, $m2$ are the medians for group 1 and 2 respectively. (Mann-Whitney U test; $+$: $p > 0.01$, $*$: $0.01 > p > 0.001$, $**$: $0.001 > p > 0.0001$, $***$: $p < 0.0001$, green represents a significant difference and a relevant effect size, yellow indicates a significant difference without a relevant effect size and red indicates a non-significant result). The groups of hypotheses adjusted for multiple correction together are marked by unique subscript numbers.

| parameter | GRHL2 | OVOL | NRF2 | GRHL2-KD |
|---|---|---|---|---|
| monostable (e)[1] | 6.6 ($***$) | 1.5 ($***$) | 1.7 ($***$) | 2.2 ($***$) |
| monostable (he)[2] | 8.4 ($***$) | 2.3 ($***$) | 2.0 ($***$) | 3.1 ($***$) |
| monostable (hm)[3] | 0.12 ($***$) | 0.51 ($**$) | 0.58 ($**$) | 0.3 ($***$) |
| monostable (m)[4] | 0.16 ($***$) | 0.53 ($***$) | 0.47 ($***$) | 0.31 ($***$) |
| p1 (e)[1] | 0.67 ($***$) | 1.1 ($*$) | 0.69 ($***$) | 1.1 ($**$) |
| p1 (he)[2] | 1.1 ($***$) | 0.94 ($*$) | 1.2 ($***$) | 0.95 (+) |
| p1 (hm)[3] | 0.99 (+) | 1.0 (+) | 0.94 (+) | 1.0 (+) |
| p1 (m)[4] | 0.92 ($*$) | 0.97 (+) | 1.0 (+) | 0.95 (+) |
| p2 (e)[1] | 1.0 (+) | 1.2 ($***$) | 0.8 ($**$) | 1.1 ($**$) |
| p2 (he)[2] | 2.6 ($***$) | 1.4 ($***$) | 1.5 ($***$) | 1.3 ($***$) |
| p2 (hm)[3] | 0.64 ($***$) | 0.75 ($***$) | 0.85 ($**$) | 0.8 ($***$) |
| p2 (m)[4] | 0.88 ($*$) | 0.87 ($***$) | 1.0 (+) | 0.99 (+) |

**Table S8:** Statistical Testing results for the PSF circuits with the proportions calculated for all the replicates of the de10 circuit forming one comparison group while the proportions of all the replicates of the oe10 circuit forming the other comparison group. The values indicate the ratio of the mean of the oe10 group and the mean of the de10 group as a measure of the effect size. (Welch's t-test; +: $p > 0.01$, $*$: $0.01 > p > 0.001$, $**$: $0.001 > p > 0.0001$, $***$: $p < 0.0001$, green represents a significant difference and a relevant effect size, yellow indicates a significant difference without a relevant effect size and red indicates a non-significant result). The groups of hypotheses adjusted for multiple correction together are marked by unique subscript numbers.)
